# A single-domain expansin-like protein from *Gloeophyllum trabeum* able to cleave xylan

**DOI:** 10.1101/2025.10.08.681111

**Authors:** Ignacio Delgado Santamaría, Heidi Østby, Vincent G. H. Eijsink, Anikó Várnai

## Abstract

Expansin-related proteins (ERPs) are a broad group of plant cell wall-loosening proteins and are considered non-catalytic as, to date, no cell wall-derived products have been observed as a result of catalysis, despite the presence of a domain that resembles the catalytic domains of GH45 endoglucanases. Here, we report catalytic activity for a single-domain ERP, *Gt*EXPN_133317, from the brown-rot fungus *Gloeophyllum trabeum*, which is highly expressed in the early phase of spruce colonization. We demonstrate enzyme-dependent formation of xylan-derived products, such as glucuronylated xylo-oligosaccharides, using high-performance anion exchange chromatography with pulsed amperometric detection. Structure-based multiple sequence alignment of ERPs with GH45 endoglucanases showed that, next to a single conserved aspartate (Asp87 in *Gt*EXPN_133317) present in all ERPs and GH45s, fungal ERPs contain a second conserved acidic residue (Asp25 in *Gt*EXPN_133317). Mutation of these two conserved amino acids, Asp87 and Asp25, led to a nearly complete loss of xylanolytic activity. While these findings do not exclude the possibility of a non-catalytic plant cell wall-loosening mechanism, they show that ERPs likely have other modes of action besides what the current paradigm states.

**Significance statement:** Protein-mediated plant cell wall-loosening is a natural process that enables plant growth and improves cell wall accessibility for microorganisms. Expansin-related proteins (ERPs) are key to this process, but their mechanism is not fully understood. Traditionally, ERPs are seen as non-catalytic, disrupting non-covalent bonds holding the cellulose network together. We have investigated an expansin-like protein from a fungal saprotroph and demonstrate its catalytic activity by showing product formation from glucuronoxylan and identifying key residues associated with this activity. This activity, although so far only shown for one protein, challenges the current paradigm that ERPs are non-catalytic, shedding new light on plant cell wall architecture and dynamics as well as on the potential roles of ERPs in plant–pathogen interactions.

## Main Text

The plant cell wall (PCW) is the main source of plant biomass, often referred to as lignocellulosic biomass, the most abundant renewable raw material on Earth. The carbohydrate fraction of lignocellulose mainly consists of cellulose, hemicellulose and pectin. The ratio of these components varies between plant species, their organs and tissues, and even individual cells, as well as between different PCW layers (1, 2). In order to colonize live plants or dead plant biomass, microorganisms secrete a wide cocktail of cell wall-modifying proteins and enzymes upon their entry through the PCW (3). Importantly, the PCW is an extremely recalcitrant, compact co-polymeric structure. It has been estimated that the average pore size in PCWs is about 35-52 Å (4), which means that proteins with a molecular mass above 40-50 kDa cannot penetrate easily.

Expansins are small (ca. 225-300 amino acids), compact, torpedo-shaped proteins consisting of an N-terminal six-stranded double-psi beta-barrel (DPBB) domain (ca. 115-150 amino acids) and a C-terminal carbohydrate-binding module (CBM; ca. 100 amino acids) belonging to the CBM63 family in the Carbohydrate-Active EnZymes database (5), connected by a short (ca. 4-amino acid long) linker (6-8). The DPBB domain is thought to enable expansins and expansin-like proteins to loosen cell walls (7). The CBM63 is unique for expansins and likely binds both cellulose and other PCW polysaccharides through two different binding surfaces (7, 9). Canonical expansins were originally discovered in plants (10) and are classified into two major families, α-expansin (EXPA) and β-expansin (EXPB), and into two minor families, expansin-like A (EXLA) and expansin-like B (EXLB) (6, 11). It is well established that EXPAs are mediators of acid-induced cell wall (CW) creep, which is the slow, irreversible, and time-dependent extension of a growing CW stretched at a constant force (12, 13). While the functions of many plant expansins remain unresolved at a molecular level, it has been shown that they are crucial for many plant processes, such as root, stem, and leaf growth, germination, reproduction, fruit/grain number and size, and resistance to abiotic and biotic stresses (14-16).

Expansin-related proteins (ERPs) are also secreted by bacteria and fungi (17). Of these, microbial expansins have the canonical DPBB–CBM63 structure and are designated as EXLX (18), the most studied being *Bs*EXLX1 from *Bacillus subtilis* (19). Microbial ERPs with non-canonical structure include loosenin-like proteins (20, 21), cerato-platanins (22, 23), and EXPNs (24, 25), which are single-domain DPBB proteins designated as ‘expansin-like proteins’ (17). ERPs also include swollenins (SWOs), which contain a canonical DPBB–CBM63 structure supplemented with an N-terminal CBM1 domain, joined by a long, unique linker (26). Microbial ERPs have been shown to promote virulence and plant colonization in a number of plant pathogens (17, 19, 27).

ERPs are considered non-catalytic proteins, which means that they are thought to act on non-covalent bonds in the PCW, such as hydrogen bonds, rather than by modifying chemical bonds (6, 9). This is remarkable considering the fact that their DPBB domains resemble the catalytic domains of endoglucanases belonging to the glycoside hydrolase (GH) family GH45 of the Carbohydrate-Active enZymes database (5), with the ability to cleave cellulose and other β-glucans (28, 29). Catalytically active GH45 enzymes contain an aspartic acid that is conserved in all DPBB proteins, including ERPs, and that acts as catalytic acid (1, 9). The position of the catalytic base varies between subfamilies of GH45 endoglucanases (30-34). Such a catalytic base has so far not been identified in ERPs (9, 35), which may help explaining why, despite numerous attempts to demonstrate catalytic activity, the current paradigm supports a non-catalytic mode of action (36-40). Regarding PCW components targeted by ERPs, in addition to cellulose, there is strong evidence for interactions with xyloglucan, which is prominent in growing PCWs (41-43). Binding to a range of other PCW components, including xylan, mannan, pectin, lignin, and mixed-linkage β-glucans has been suggested (7, 39, 44-50). However, whether binding to these polysaccharides occurs through the DPBB or the CBM63 needs further investigation.

Herein, we present the biochemical characterization of a single-domain EXPN protein, denoted as *Gt*EXPN_133317, from the fungal saprophyte *Gloeophyllum trabeum* (51, 52). Using high-performance anion exchange chromatography (HPAEC) and mass spectrometry (MS) analyses, we demonstrate that *Gt*EXPN_133317 exhibits catalytic activity towards xylan, producing linear and branched xylan fragments of varying length. Structure-based multiple sequence alignment of plant, bacterial, and fungal ERPs with GH45 endoglucanases indicates the potential occurrence of a hitherto unrecognized catalytic base in ERPs. The importance of this putative catalytic base, and of the catalytic acid, was then confirmed by mutagenesis studies. The discovery of a hemicellulolytic hydrolytic activity in ERPs sheds new light on how expansins and expansin-like proteins may function in nature.

## Results and discussion

### *Gt*EXPN_133317, a single-domain fungal expansin-like protein from the brown-rot fungus *G. trabeum*, and its relation to GH45s

EXPNs are single-domain proteins lacking the CBM63 typically found in plant and microbial expansins, and are widely found in fungal genomes (17). The genome of *G. trabeum* (51) encodes nine such EXPNs (**Table S1**). Transcriptome analyses of *G. trabeum* grown on aspen wood wafers (53) and on glucose, cellulose, and Japanese cedar (54) have shown that the genes of all nine *Gt*EXPNs are transcribed. During growth on aspen wood wafers and glucose, *Gt*EXPN_133317 transcripts were detected at levels that were one to two orders of magnitude higher compared to the other seven *Gt*EXPNs. In contrast, *Gt*EXPN_130898 was the most abundant transcript during growth on cellulose and Japanese cedar (**Table S2**). Secretome analyses (**Table S3**) have shown that of the nine *Gt*EXPNs, *Gt*EXPN_133317 and *Gt*EXPN_44870 are dominant in the early secretome when *G. trabeum* grows on spruce or aspen wood wafers (55, 56). *Gt*EXPN_44870 and *Gt*EXPN_130898 share 62% sequence identity, while *Gt*EXPN_133317 shares 47% and 46% with *Gt*EXPN_44870 and *Gt*EXPN_130898, respectively (**Table S1**). Because of its abundance in both secretomes and transcriptomes as well as its occurrence in the early secretome, we selected *Gt*EXPN_133317 for further characterization.

The structural model of *Gt*EXPN_133317 suggested by AlphaFold3 (57) confirms a DPBB-like structure (**Figure 1A-B**) that is similar to the experimentally determined structures of the N-terminal DPBB domains of plant (*Zm*EXPB1 from maize [PDB: 2HCZ] (48), *Gh*EXLA1 from cotton [PDB: 7XC8] and *Phlp*1 from Timothy grass [PDB: 1N10]), bacterial (*Bs*EXLX1 from *Bacillus subtilis* [PDB: 4FG4] (19) and *Cm*EXLX from *Clavibacter michiganensis* [PDB: 4JCW]) and fungal (*Tl*EXLX from *Talaromyces leycettanus* [PDB: 7WVR] (58)) expansins and to the single-domain fungal loosenin-like protein *Pca*LOOL12 from *Phanerochaete carnosa* [PDB: 9CE9] (59). The model shows a flexible disordered region at the N-terminus (Ser1-Thr16; **Figure 1B** and **Figure S1)**, as also observed for single-domain fungal loosenin-like proteins (59). While the predicted confidence of the model was low for this N-terminal tail, the predicted confidence was uniformly high for the rest of the protein, which adopts the DPBB fold (**Figure S1**).

**Figure 1.**
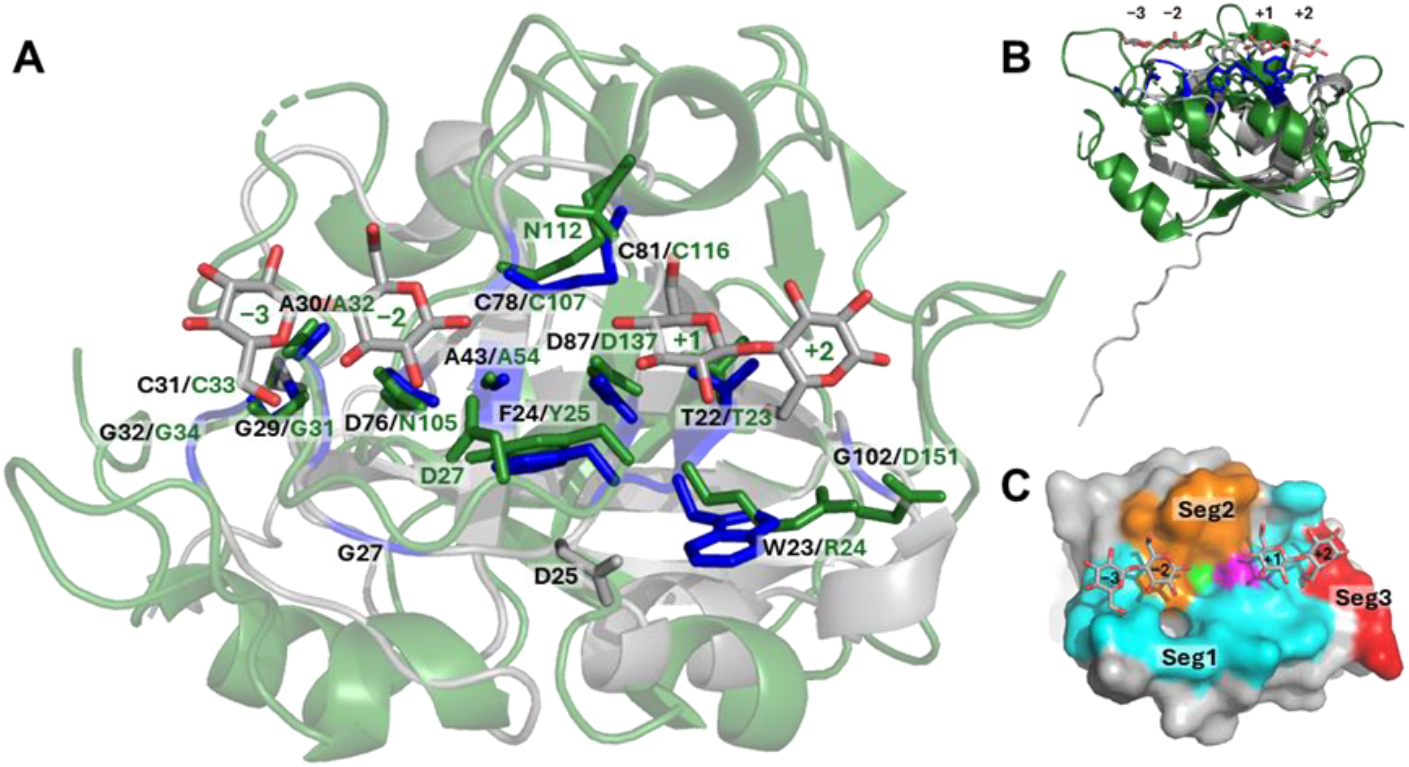
Potential substrate-binding and catalytic residues of fungal EXPNs. **A)** Structural alignment of *Gt*EXPN_133317 (grey, AlphaFold3 model) and *Ac*Cel45A in complex with cellobiose [5XC8] (green). Selected amino acid residues conserved in brown-rot fungal EXPNs (**Figure S5**) are labelled in black and shown in blue for *Gt*EXPN_133317. Corresponding residues and the putative catalytic base (Asn112) in *Ac*Cel45A are labelled and shown in green. The sugar-binding subsites of *Ac*Cel45A (30) are labelled for reference. **B)** Sideview of panel **A**. Note the N-terminal disordered region in *Gt*EXPN_133317. **C)** Segments of the substrate-binding surface and the catalytic centre of *Gt*EXPN_133317, colored according to the annotation in **Figure 2**. The conserved Asp87 is shown in magenta; the alanine (Ala43) forming the bottom of a pocket next to the aspartate is shown in green. The picture also shows the cellobiose from the structure of *Ac*Cel45A [5XC8] based on the structural alignment shown in panel **A**.

The DPBB domain in ERPs and GH45s contains six β-strands that are connected by loops that vary in length (60) (**Figure 2**). Variation in some of these loops translates into variations in the nature and size of the substrate-binding surface (**Figures S2-S4**). The core DPBB fold of *Gt*EXPN_133317 aligns well with the structure of GH45s from subfamilies B (*Ac*Cel45A [PDB: 5XC8] and *Me*Cel45A [PDB: 1WC2]; **Figure 1A**) and one member of subfamily C (*Gt*Cel45A [PDB: 8BZQ]) but less so with those from subfamily A (e.g., *Hi*Cel45A [PDB: 3ENG]). Here, we have defined the regions of the protein that make up the substrate-binding surface around the fully conserved aspartate that serves as a catalytic acid in GH45s. For all ERPs, the potential substrate-binding surface is formed by three segments (Seg1-Seg3) and two amino acids, Ala43 and Asp87 in *Gt*EXPN_133317 (**Figures 1C & 2**). Surface residues on these segments vary between GH45s and ERPs, but the ERPs show clear conservation patterns (35). A comparison of EXPNs from nine brown-rot strains revealed twelve conserved residues in the substrate binding surface (**Figure S5**), including Thr22, which is followed by two conserved aromatic residues (Trp23 and Phe24 in *Gt*EXPN_133317) and an aspartate (Asp25) that is discussed below. Generally, the loops connecting the β-strands are longer in GH45s compared to ERPs, which gives GH45s a more extended substrate-binding surface (**Figures 2 & S2**) and the ability to form clefts or even a tunnel to accommodate a polysaccharide substrate (**Figures S3 & S4**). In contrast, ERPs have smaller and flatter binding surfaces (**Figure S3A-C & S4A-C**).

**Figure 2.**
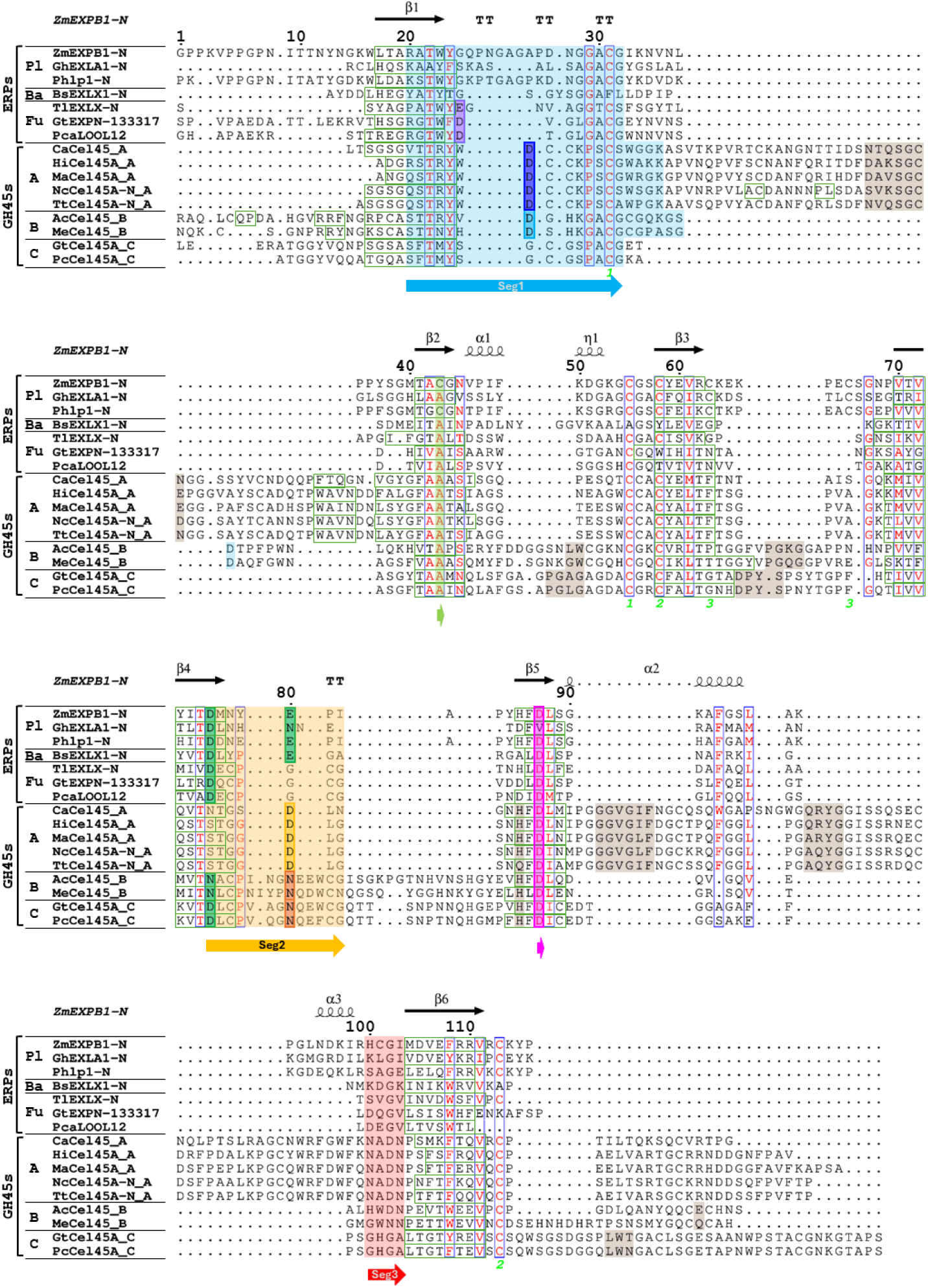
Structure-based multiple sequence alignment of *Gt*EXPN_133317 with characterized expansins and GH45s. PDB structures: Plant (Pl) expansins: *Zm*EXPB1 [2HCZ], *Gh*EXLA1 [7XC8], *Phlp*1 [1N10]; Bacterial (Ba) expansin: *Bs*EXLX1 [4FG4]; Fungal (Fu) expansin-related proteins: expansin *Tl*EXLX [7WVR], EXPN *Gt*EXPN_133317 [predicted structure, this study], loosenin-like protein *Pca*LOOL12 [9CE9]; GH45 subfamily A: *Ca*Cel45 [5H4U], *Hi*Cel45A [3ENG], *Ma*Cel45A [1OA7], *Nc*Cel45A [6MVJ], *Tt*Cel45A [5GLX]; subfamily B: *Ac*Cel45 [5XC8], *Me*Cel45 [1WC2]; subfamily C: *Gt*Cel45A [8BZQ]; *Pc*Cel45A [3X2M]. Structural elements indicated above the alignment are based on the *Zm*EXPB1 structure [PDB: 2HCZ]. β-Strands are marked with green boxes; the β-strands (β1-β6) forming the β barrel are indicated above the sequence. Residues are numbered based on the residues in *Gt*EXPN_133317. The segments making up the substrate-binding surface, as defined in this study, appear on colored background, and the segments present in ERPs (Seg1-Seg3 and two amino acids) are labelled with colored arrows below the alignment. Residues with known or plausible catalytic functions in GH45s appear in colored boxes, as follows: magenta, catalytic acid (Asp); dark blue, catalytic base in subfamily A GH45s (Asp); dark orange, catalytic base in subfamily B and C GH45s (Asn); light orange, important substrate-coordinating residue in subfamily A GH45s (Asp); light blue, important substrate-coordinating residue in subfamily B GH45s (Asp). Residues identified in the present study as being important for catalytic activity of *Gt*EXPN_133317 appear in magenta and purple boxes; other conserved residues for which the possibility to contribute to catalysis in ERPs is discussed in this study appear in green boxes. Green numbers under the sequence indicate cysteine bridges in *Zm*EXPB1 [PDB: 2HCZ].

The structure-based alignment of the DPBB domains of ERPs and GH45 (**Figure 2**) shows that the catalytic acid that is associated with cellulolytic activity in GH45s is fully conserved in ERPs (Asp87 in *Gt*EXPN_133317). In inverting GHs, such as GH45 (61), one would expect to find a conserved second acid residue acting as a catalytic base at some 10 Å from the catalytic acid (1). Interestingly, the location and nature of this base is not clear for all studied GH45s and seems to vary among the enzymes. Subfamily A GH45s (GH45_As) contain an aspartate functioning as the catalytic base (32), whereas asparagine residues located elsewhere in the protein have been proposed to play that role in Subfamily B and C GH45s (30, 31, 33); see **Figure 2** for details).

While the catalytic acid of GH45s is conserved, full GH45-like catalytic arrangements do not occur in ERPs. Resemblances do occur, however, in some ERPs. For example, several ERPs, but not *Gt*EXPN_133317, contain a glutamate or asparagine at the position of the putative catalytic base in GH45_Bs and GH45_Cs (**Figures 2 & 3**). The corresponding Glu75 in *Bs*EXLX1, for which no catalytic activity has been detected, has been found important but not essential for creep activity (7). In fungal ERPs, including *Gt*EXPN_133317, this residue is a glycine, and there are no nearby residues that can play catalytic roles. *Gt*EXPN_133317 contains Asp25, at a similar but still conspicuously different location as the conserved aspartate that has been implied in catalytic activity of GH45_As and GH45_Bs (30, 32) (**Figure 3)**, at 9.6 Å from Asp87 **(Figure S6)**. Furthermore, *Gt*EXPN_133317 contains an aspartate (Asp76) that is conserved in fungal and bacterial ERPs (35), with a predicted distance of 6.7 Å from Asp87 (**Figure S6**), which aligns with a conserved aspartate in GH45_Cs that has been proposed to play a role during catalysis (31).

**Figure 3.**
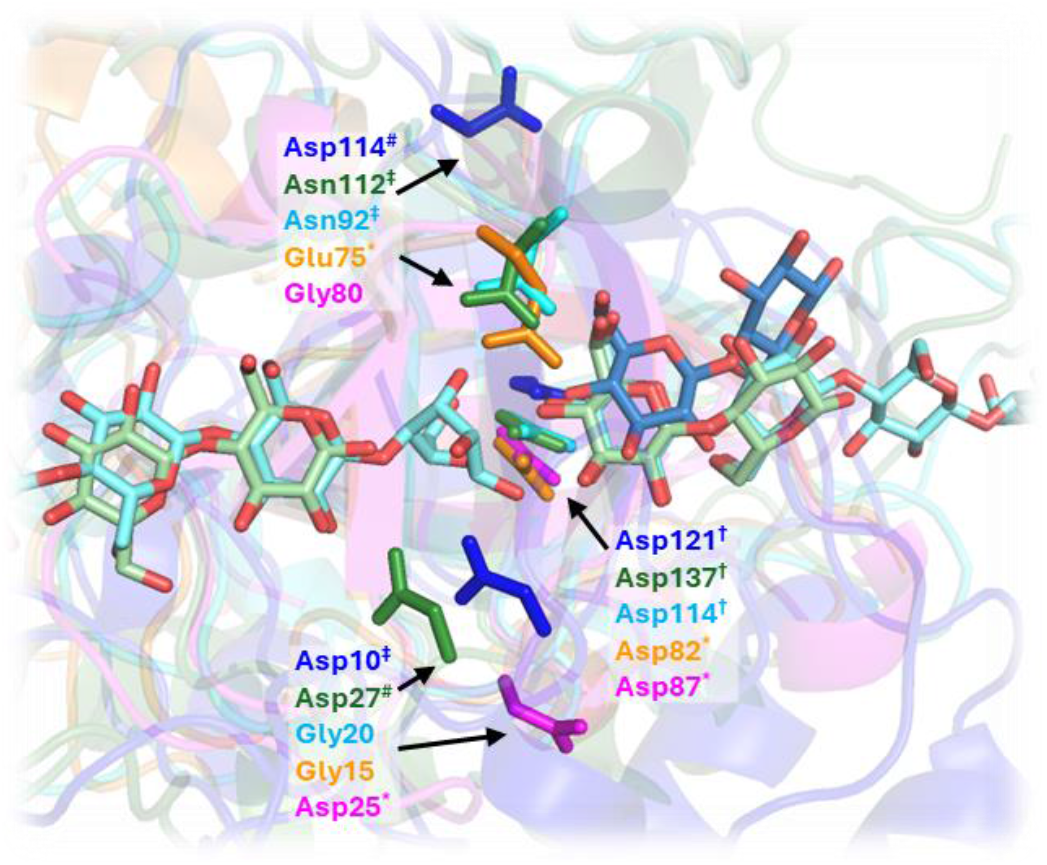
Conserved amino acid residues in the catalytic centers of ERPs and GH45_B and GH45_C endoglucanases. ERPs: *Bs*EXLX1 [4FG2], orange; *Gt*EXPN_133317, magenta. Endoglucanases: *Hi*Cel45A_A with cellobiose [3ENG], dark blue; *Ac*Cel45A_B with cellobiose [5XC8], dark green; *Pc*Cel45A_C with cellopentaose [3X2M], light blue. For the GH45 endoglucanases, the experimentally determined catalytic acids and bases in *Hi*Cel45A_A (32), *Ac*Cel45A_B (30) and *Pc*Cel45A_C (33) are indicated with † and ‡, respectively. In some GH45s, a third, assisting residue has been identified and is marked with #. Mutating one of these three residues results in complete or severe loss of protein activity. Note that the position of the catalytic base varies between these GH45s. Two residues that are essential for plant CW-loosening activity in *Bs*EXLX1 (7) and xylanolytic activity in *Gt*EXPN_133317 (this study) are marked with *. For both proteins one of these residues aligns with the catalytic acid in GH45s (Asp87 in *Gt*EXPN_133317, Asp82 in *Bs*EXLX1) whereas the position of the second residue differs (Asp25 in *Gt*EXPN_133317, Glu75 in *Bs*EXLX1).

### *Gt*EXPN_133317 has xylanolytic activity

To date, expansins are generally considered non-catalytic proteins (15, 17). To challenge this dogma, we set out to characterize *Gt*EXPN_133317. The gene coding for this protein was cloned and expressed in *Pichia pastoris* KM71H, recently renamed *Komagataella phaffii*, (*Gt*EXPN_133317^*Pp*^) and *Escherichia coli* BL21(DE3) (*Gt*EXPN_133317^*Ec*^), and the proteins were purified by anion exchange and size exclusion chromatography. SDS-PAGE analysis indicated a difference in the molecular masses of *Gt*EXPN_133317 produced in the two different hosts (**Figure S7**). MALDI-ToF MS analysis of the purified proteins (**Figure S8**) confirmed that *Gt*EXPN_133317^*Ec*^ was correctly produced, with the pelB leader sequence being correctly cleaved off and without proteolytic damage. In contrast, the N terminus (Ser1–Arg14), and to a lesser extent also the C terminus (Phe115–Pro117), of *Gt*EXPN_133317^*Pp*^ underwent proteolytic cleavage, yielding several smaller, partially truncated proteins (**Table S4** and **Figure S9**). MALDI-ToF MS analysis indicated no post-translational modifications (**Table S4** and **Figure S9**), which was confirmed with peptide mapping with LC-MS^*2*^ analysis of *Gt*EXPN_133317^*Pp*^ (**Dataset S1**). The proteolytic cleavage likely led to reduced stability as indicated by a lower apparent melting temperature (T^*m*(app)^) for *Gt*EXPN_133317^*Pp*^ compared to *Gt*EXPN_133317^*Ec*^ (**Figure S10**).

We evaluated the activity of *Gt*EXPN_133317 on PCW-derived polysaccharides by assessing changes in rheology or turbidity. These methods required very high xylan concentrations, and we were not able to obtain conclusive results. Conclusive results were, however, obtained when analyzing soluble products using high performance anion exchange chromatography (HPAEC-PAD) and mass spectrometry (MALDI-ToF MS). Since expansins are generally considered to act on (and loosen) copolymeric substrates in the PCW, we performed the reactions using a mixture of beechwood xylan (4-*O*-methylglucuronylated xylan) and amorphous cellulose (PASC) as substrate, using reactions with PASC or beechwood xylan alone as controls. Combining PASC with xylan has previously enabled the discovery of xylanolytic activity for a group of AA9 lytic polysaccharide monooxygenases, which only act on xylan that is associated to cellulose (62, 63).

HPAEC-PAD revealed that both protein variants (*Gt*EXPN_133317^*Pp*^ and *Gt*EXPN_133317^*Ec*^) released a broad range of xylan fragments when acting on a mixture of beechwood glucuronoxylan and PASC (**Figure 4**). Purified *Gt*EXPN_133317^*Pp*^ also exhibited weak cellulolytic activity on PASC (**Figure 4A**). However, this cellulolytic activity can be attributed to an endoglucanase background that was not removed entirely during purification, as the culture broth of the non-transformed *P. pastoris* expression host also exhibited this activity (**Figure S11**). Activity on glucuronoxylan was not detected in such broth. Importantly, cellulolytic background activity was not detected for purified *E. coli*-produced *Gt*EXPN_133317^*Ec*^ (**Figure 4B**). Additional activity assays showed that *Gt*EXPN_133317 is also active on beechwood glucuronoxylan alone (**Figure S12**). Heat-inactivation of *Gt*EXPN_133317^*Ec*^ abolished product formation (**Figure S13**).

**Figure 4.**
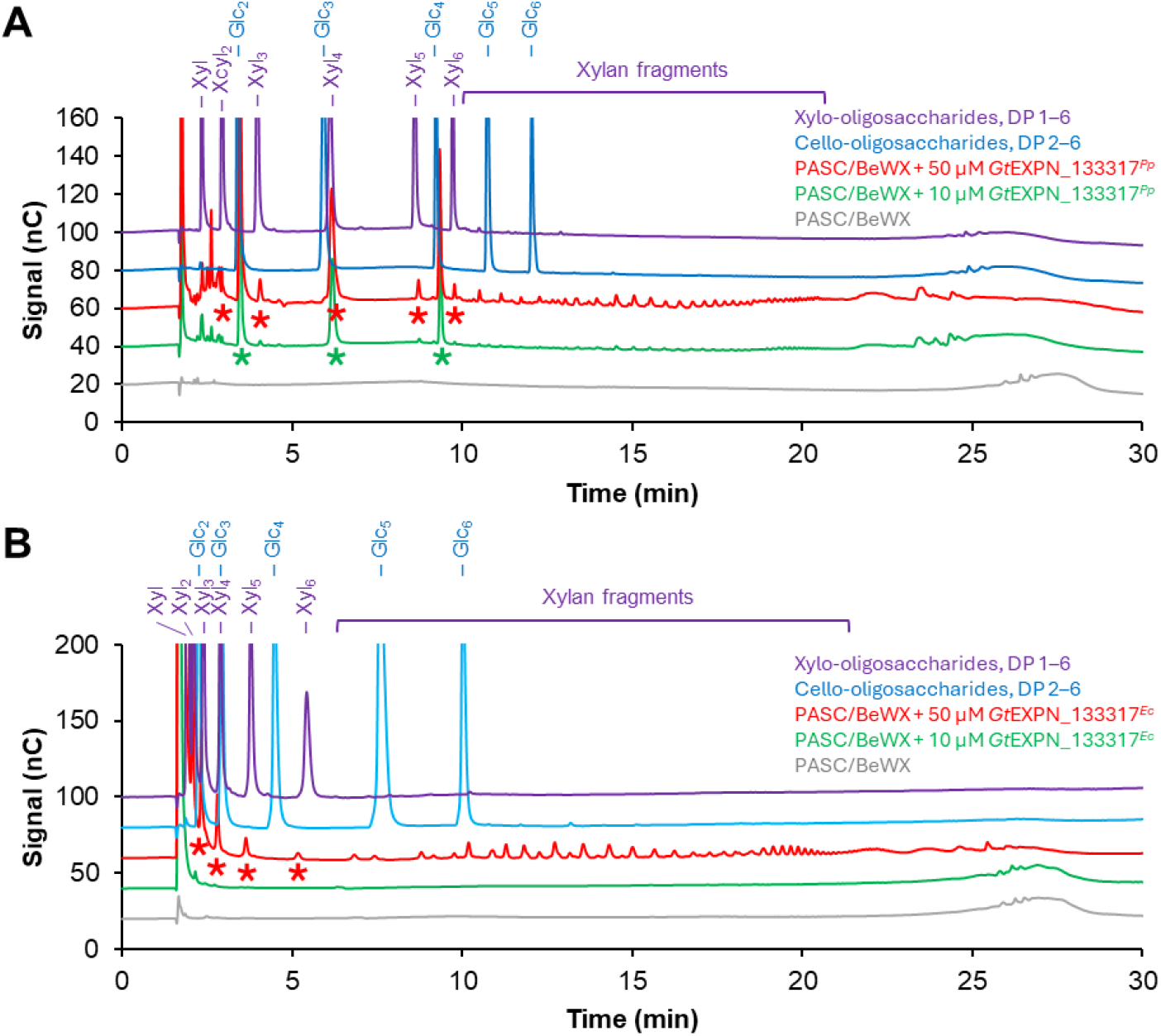
HPAEC-PAD analysis of products generated in a reaction with *Gt*EXPN_133317 and a 1:1 (w/w) mixture of PASC and beechwood xylan. **A)** *Gt*EXPN_133317^*Pp*^ **B)** *Gt*EXPN_133317^*Ec*^. In the reactions, 10 µM (green line) or 50 µM (red line) *Gt*EXPN_133317 was incubated with 0.1% (w/v) beechwood xylan + 0.1% (w/v) PASC at 30°C, pH 5.0, for 48 h with horizontal shaking at 200 rpm. The identified cello-oligosaccharides (with DP 2–4) are marked with green asterisks; the identified linear xylo-oligosaccharides (with DP 2–6) are marked with red asterisks. Chromatograms for control reactions without enzyme are shown with grey lines. All reactions were performed in triplicates, with similar results; a typical chromatogram is shown. Cello-oligosaccharides with a degree of polymerization (DP) 2–6 (blue line) and xylo-oligosaccharides with DP 1–6 (purple line), each at 0.025 mg/mL concentration, were used as standard. There is a shift in the retention times between panels A and B due to the sensitivity of HPAEC separation to CO_2_ penetrating into the eluents and affecting retention of the analytes (81). Chromatograms for enzyme reactions with only PASC were very similar to the green chromatograms in panels A and B, conforming background glucanase activity for *Gt*EXPN_133317^*Pp*^ and the absence of such activity for *Gt*EXPN_133317^*Ec*^. Chromatograms for enzyme reactions with only beechwood xylan are shown in **Figure S12** and are discussed in the main text.

Xylanolytic activity was confirmed using MALDI-ToF MS (**Figure 5**), which showed the formation of xylan-derived fragments of varying lengths, including linear (unsubstituted) fragments (Xyl_n_; up to *n*=17), singly glucuronylated fragments (Xyl_*n*_MeGlcA; *n*=4–17) and doubly glucuronylated fragments (Xyl_*n*_MeGlcA_2_; *n*=8–19). The *m/z* values of the detected compounds corresponded to those of hydrolysis products, and the degree of product glucuronylation corresponded to that of the employed hardwood-derived xylan substrate, which is expected to carry this modification primarily at every sixth xylosyl residue (64, 65). It is noteworthy that, after taking a closer look at the spectrum, we detected species that could correspond to dehydrated species [marked with ‘(−H_2_O)’], albeit at low levels (**Figure S14**). While our MALDI-ToF MS data suggest a hydrolytic type of action, further studies may be needed to confirm the nature of the reaction products.

**Figure 5.**
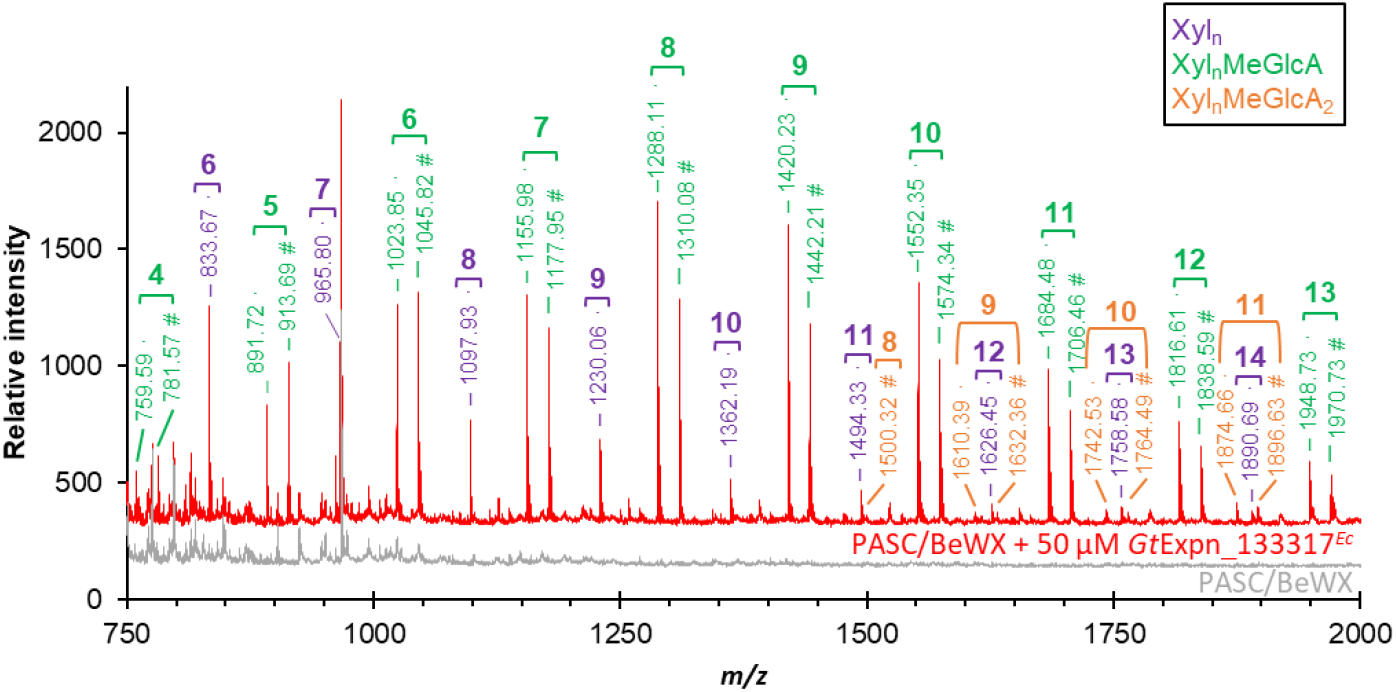
MALDI-ToF MS analysis of the products generated in a reaction of *Gt*EXPN_133317^*Ec*^ with a 1:1 (w/w) mixture of PASC and beechwood xylan. Products from the catalytic action of *Gt*EXPN_133317^*Ec*^ (red line) include non-substituted (purple labels; Xyl_*n*_) and singly (green labels; Xyl_*n*_MeGlcA) or doubly (orange labels; Xyl_*n*_MeGlcA_2_) glucuronylated xylan fragments of varying lengths, where *n* is the number of xylosyl units in the xylan fragment backbone, which is indicated above square brackets for each product. The *m/z* ratios of Na^+^-adducts of the products are indicated and are followed by ‘·’ for single sodium adducts or by ‘#’ for the Na^+^-salt of the corresponding compounds for singly (green) and doubly (orange) glucuronylated xylan fragments. A control reaction without *Gt*EXPN_133317^*Ec*^ is shown with a grey line. A close-up of the products in the *m/z* range of 1520–1670 is shown in **Figure S14**. The theoretical *m/z* values of glucuronoxylan-derived compounds are listed in **Table S5**.

Xylanolytic activity was also assessed using the linear oligomers xylopentaose and xylohexaose as substrates. HPAEC-PAD and MS analysis of product formation revealed that *Gt*EXPN_133317 is able to cleave xylopentaose and xylohexaose (**Figures S15-S16**), generating xylobiose and xylotriose, or xylotetraose, xylotriose and xylobiose, respectively. The rate of these reactions was extremely low (on the order of a few per hour), which indicates that these oligomeric substrates are not true substrates for *Gt*EXPN_133317.

Xylan is an important constituent of the PCW where it has abundant contacts with cellulose microfibrils and lignin (66, 67). Correspondingly, PCW-associated microorganisms have developed a diverse set of enzymatic machinery to modify or depolymerize xylan (68), and this machinery may, as we show here, include ERPs. Due to the lack of standards, it was not possible to determine the rate of the reaction with beechwood xylan. It is clear though, that the rate is low and that substrate conversion is limited, since we could only detect products by HPLC and mass spectrometry and not by other assays such as measuring reducing end formation or monitoring changes in viscosity or turbidity (results not shown). Low activity aligns well with the notion that ERPs have evolved to cleave key biomechanical junctions between cell wall polymers (42, 69). Thus, *Gt*EXPN_133317 may have evolved to selectively cleave a limited number of inaccessible junction sites, perhaps at a deliberately slow rate compatible with plant growth, thereby loosening interfaces between xylan and other plant cell wall components.

Activity towards xylan has been suggested previously for *Tr*SWO, a swollenin from *T. reesei*, but this activity was attributed to the release of oligosaccharides that pre-existed within the substrate (70). For fungal-expressed *Tr*SWO, low activity on β-(1→3),(1→4)-glucan and even weaker (orders of magnitude lower) activity on birchwood xylan have also been suggested (70, 71). However, due to the endogenous glucanases and xylanases produced by the fungal expression hosts, it has not been possible to convincingly demonstrate that these activities are not attributable to background activity. Phylogenetic analysis of the DPBB domains of 125 ERPs (**Figure S17**) shows that *Gt*EXPN_133317 is part of a clearly separate clade of single-domain EXPNs containing *Pca*LOOL12. Recently, *Pca*LOOL12 has been shown to impact the arrangement of cellulose microfibrils in spruce (softwood) and Eucalyptus (hardwood) (59), which could potentially be due to a hitherto undiscovered action on recalcitrant xylan coating cellulose microfibrils in both types of woody biomass (65). This, however, remains to be confirmed.

### Mutagenesis reveals key catalytic residues in *Gt*EXPN_133317

In order to prove that the observed xylanolytic activity of *Gt*EXPN_133317 is indeed due to this protein (and not contaminations) and to identify catalytically important residues, the aspartate residues discussed above (Asp87, Asp76, and Asp25) were mutated, generating the D87A and D25N mutants. Proteins in which Asp76 was mutated (D76A and D76N) were expressed only in very low quantities, and we could not produce enough protein for activity tests. AlphaFold3 predicted that these mutants would fold correctly, albeit with a change in hydrogen bonds involving residue 76 (**Figure S18**).

HPAEC-PAD analysis confirmed that the D87A and D25N mutations both were detrimental to the xylanolytic activity of *Gt*EXPN_133317 (**Figure 6**). Mutation of the Asp87 to alanine (D87A) completely eliminated the activity of *Gt*EXPN_133317 on beechwood xylan, as one would expect based on previous mutational studies of GH45s, where this Asp is thought to act as catalytic acid (9, 30, 32, 33). Mutation of Asp25 to asparagine led to a substantial reduction in activity, with small amounts of products still being detected. This indicates that Asp25 is also essential for the xylanolytic activity. Control experiments showed that both mutants had thermal stabilities higher than wild-type *Gt*EXPN_133317^*Pp*^ (**Figure S19**), indicating that the lack of observed activity was not due to misfolding or thermal inactivation.

**Figure 6.**
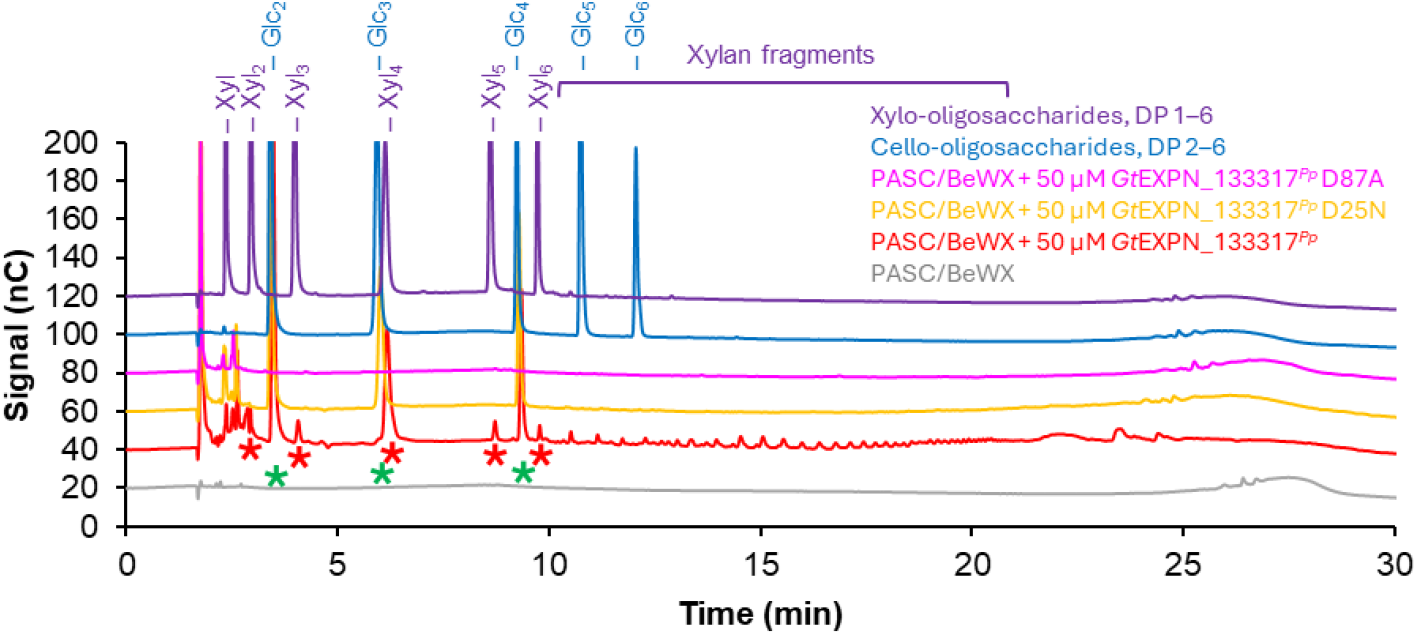
HPAEC-PAD analysis of products generated in a reaction with *Gt*EXPN_133317^*Pp*^ and mutants and a 1:1 (w/w) mixture of PASC and beechwood xylan. In the reactions, 50 µM *Gt*EXPN_133317^*Pp*^ (red line), *Gt*EXPN_133317^*Pp*^ D25N (orange line) or *Gt*EXPN_133317^*Pp*^ D87A (magenta line) was incubated with 0.1% (w/v) beechwood xylan + 0.1% (w/v) PASC at 37°C, pH 5.0, for 48 h with horizontal shaking at 200 rpm. The identified cello-oligosaccharides (with degree of polymerization, DP, 2–4) are marked with green asterisks; the identified linear xylo-oligosaccharides (with DP 2–6) are marked with red asterisks. The result for a control reaction without enzyme is shown with a grey line. All reactions were performed in duplicates, with similar results; a typical chromatogram is shown (see **Figure S26** for all the chromatograms). Cello-oligosaccharides with a DP 2–6 (blue line) and xylo-oligosaccharides with DP 1–6 (purple line), each at 0.025 mg/mL concentration, were used as standard.

It is noteworthy that the cellulolytic activity was detected for wild-type *Gt*EXPN_133317^*Pp*^ and the D25N mutant, but not for the D87A mutant (**Figure 6**), despite subjecting the *P. pastoris* broths to identical purification steps. The most likely explanation for this is a difference in the extent of removal of the endoglucanase background during purification. Indeed, SDS-PAGE analysis of the purified protein preparations showed less background proteins in the D87A sample (**Figures S7** and **S20**).

### The impact of reaction parameters on the xylanolytic activity of *Gt*EXPN_133317

We then assessed the impact of time, xylan concentration, enzyme loading, and pH on product formation. While increasing the reaction time (**Figure S21**) and enzyme concentration (**Figure 4**) led to increased product formation, increasing the xylan concentration (**Figure S22**) had no impact on product release by *Gt*EXPN_133317. Furthermore, changing the buffer from 20 mM sodium acetate (pH 5.0) to 50 mM BisTris/HCl (pH 6.5) significantly increased the xylanolytic activity of *Gt*EXPN_133317 (**Figure S23**). Changing the buffer to 25 mM MES/KOH at pH 6.5 had no significant impact on the level of product formation, whereas employing 25 mM HEPES/KOH (pH 7.5) and 25 mM Tris/HCl (pH 9.5) led to a substantial reduction in activity (**Figure S24**). Overall, with the conditions used here, the xylanolytic action of *Gt*EXPN_133317 seemed limited by the enzyme rather than by the substrate.

It is noteworthy that α-expansins are thought to induce creep in seconds at acidic pH (4-5) (10, 72, 73), while in the present study xylan-derived products were only observed after many hours of incubation. This latter timescale aligns well with the reported effects of several expansin-like proteins on cellulose and hardwood pulp integrity (47, 74, 75). Interestingly, microbial expansins have shown more neutral pH optima for their creep-inducing activity (between 5.5-9.5 for *Bs*EXLX1) (7, 15, 76), which is compatible with our observation that *Gt*EXPN_133317 is more active at pH 6.5 than at pH 5.0.

### Substrate specificity of *Gt*EXPN_133317 and other ERPs

In PCWs, xylan adopts a two-fold screw conformation when coating cellulose fibres (77, 78) and a preferential three-fold screw conformation when it is not bound to and constrained by cellulose, free in solution, or bound to lignin (66, 67, 79, 80). The xylanolytic activity of *Gt*EXPN_133317 on soluble deacetylated beechwood glucuronoxylan could indicate that *Gt*EXPN_133317 cleaves xylan with a three-fold screw conformation. To gain further insight into expansin action, the specific characteristics of xylan in natural PCWs, including its substitutions and conformation, need to be further elucidated. Literature data indicate considerable variation in terms of the PCW components with which ERPs interact. PCW loosening by plant and bacterial expansins is thought to occur not only at the interface between cellulose fibres but also at the interface between cellulose and xyloglucan (41-43). On the other hand, some microbial expansins preferentially bind to xylan and alter cellulose networks in cellulose pulps (47), while the fungal swollenin *Tr*SWO has been suggested to preferentially target mixed-linkage β-glucan (70, 71). Thus, we incubated *Gt*EXPN_133317 with Avicel microcrystalline cellulose, Whatman No. 1 filter paper, xyloglucan from tamarind, mannan from ivory nut, mixed-linkage glucan from barley, and polygalacturonic acid. Despite an extended incubation period of 48 h, no product formation was observed beyond what was anticipated from the cellulolytic background in the protein preparation (**Figure S25**). Of note, the lack of activity on these isolated, soluble PCW components does not exclude potential catalytic activity on these substrates when part of complex co-polymeric PCW structures.

### Concluding remarks

To the best of our knowledge, this study provides the first convincing example of a polysaccharide-degrading activity in ERPs. Slow cleavage and, in the case of plant growth, re-assembly of elements in the co-polymeric PCW provide a logical explanation for expansin function. It remains to be seen whether the activity detected here is universal for ERPs. The sequence conservation patterns discussed above suggest that similar activities exist in at least a subfraction of ERPs.

Several obstacles have been hindering the quest for catalytic activity in ERPs, including the presence of interfering carbohydrate-active background enzymes originating from expression hosts, inappropriate reaction setups (suboptimal reaction conditions; lack of suitable controls), the use of inadequate or insensitive methods for product detection, and, to some extent, our assumption that ERPs are non-catalytic. The discovery of xylanolytic activity in an ERP can be attributed to the high sensitivity of HPAEC-PAD and mass spectrometry used for product detection. These methods have rarely been used in the search of expansin-catalyzed reactions. Notably, these methods are able to detect products independent of the type of cleavage and are thus independent of whether the cleavage of the glycosidic bond by ERPs generates a reducing end or not. While working with these proteins, we noted that they tend to stick to surfaces (hence we had to use glass rather than plastic tubes) and their rather limited stability (hence we had to use relatively low temperatures and moderate mixing). These potential pitfalls should be taken into account when looking for the catalytic activities of ERPs.

The active site structures of the different types of ERPs likely have evolved similarly to those in the GH45 subfamilies. Although a non-catalytic PCW-loosening action by ERPs cannot be excluded, the present results confirm that catalytically active ERPs do exist. Thus, ERPs likely have modes of action besides what the current paradigm states, that is, inducing creep and targeting non-covalent bonds in the PCW (15). Both protein families, GH45s and ERPs, show considerable variation in their (putative) catalytic centers, indicating that several catalytic mechanisms may exist. Since the term ‘expansin-like proteins’ covers proteins with varying structures and sequences, and based on the present results, it is likely that that new catalytic activities will be discovered in the years to come.

## Materials and Methods

See Supporting Information.

## Acknowledgments

We thank Novo Nordisk Foundation for funding this research through an Emerging Investigator grant to Anikó Várnai [grant no. NNF-0061165]. We thank Dr. Peter Elias Kidibule for his contributions to the generation of *Gt*EXPN_133317 mutants. This work was co-funded by the Research Council of Norway through grant no. 270038 (NorBioLab).

## References

1. C. M. Payne et al., Fungal cellulases. Chemical Review 115, 1308–1448 (2015).

2. M. Pauly, K. Keegstra, Cell-wall carbohydrates and their modification as a resource for biofuels. Plant J. 54, 559–568 (2008).

3. C. P. Kubicek, T. L. Starr, N. L. Glass, Plant cell wall-degrading enzymes and their secretion in plant-pathogenic fungi. Annu. Rev. Phytopathol. 52, 427–451 (2014).

4. N. Carpita, D. Sabularse, D. Montezinos, D. P. Delmer, Determination of the pore size of cell walls of living plant cells. Science 205, 1144–1147 (1979).

5. E. Drula et al., The carbohydrate-active enzyme database: functions and literature. Nucleic Acids Res. 50, D571–D577 (2022).

6. D. J. Cosgrove, Plant expansins: diversity and interactions with plant cell walls. Curr. Opin. Plant Biol. 25, 162–172 (2015).

7. N. Georgelis, A. Tabuchi, N. Nikolaidis, D. J. Cosgrove, Structure-function analysis of the bacterial expansin EXLX1. J. Biol. Chem. 286, 16814–16823 (2011).

8. A. B. Boraston, D. N. Bolam, H. J. Gilbert, G. J. Davies, Carbohydrate-binding modules: fine-tuning polysaccharide recognition. Biochemical Journal 382, 769–781 (2004).

9. N. Georgelis, N. Nikolaidis, D. J. Cosgrove, Bacterial expansins and related proteins from the world of microbes. Appl. Microbiol. Biotechnol. 99, 3807–3823 (2015).

10. S. McQueen-Mason, D. M. Durachko, D. J. Cosgrove, Two endogenous proteins that induce cell wall extension in plants. Plant Cell 4, 1425–1433 (1992).

11. J. Sampedro, D. J. Cosgrove, The expansin superfamily. Genome Biology 6, 242 (2005).

12. D. J. Cosgrove, Catalysts of plant cell wall loosening. F1000Research 5 (2016).

13. D. J. Cosgrove, Loosening of plant cell walls by expansins. Nature 407, 321–326 (2000).

14. P. Marowa, A. Ding, Y. Kong, Expansins: roles in plant growth and potential applications in crop improvement. Plant Cell Rep. 35, 949–965 (2016).

15. D. J. Cosgrove, Plant cell wall loosening by expansins. Annual Review of Cell and Developmental Biology 40, 329–352 (2024).

16. M. Samalova, E. Gahurova, J. Hejatko, Expansin-mediated developmental and adaptive responses: a matter of cell wall biomechanics? Quantitative Plant Biology 3 (2022).

17. D. J. Cosgrove, Microbial Expansins. Annu. Rev. Microbiol. 71, 479–497 (2017).

18. H. Kende et al., Nomenclature for members of the expansin superfamily of genes and proteins. Plant Mol. Biol. 55, 311–314 (2004).

19. F. Kerff et al., Crystal structure and activity of Bacillus subtilis YoaJ (EXLX1), a bacterial expansin that promotes root colonization. Proceedings of the National Academy of Sciences 105, 16876–16881 (2008).

20. R. E. Quiroz-Castañeda, C. Martínez-Anaya, L. I. Cuervo-Soto, L. Segovia, J. L. Folch-Mallol, Loosenin, a novel protein with cellulose-disrupting activity from Bjerkandera adusta. Microbial Cell Factories 10, 8 (2011).

21. D. Dahiya et al., Fungal loosenin-like proteins boost the cellulolytic enzyme conversion of pretreated wood fiber and cellulosic pulps. Bioresour. Technol. 394, 130188 (2024).

22. L. Pazzagli et al., Purification, characterization, and amino acid sequence of cerato-platanin, a new phytotoxic protein from ceratocystis fimbriata f. sp. platani. J. Biol. Chem. 274, 24959–24964 (1999).

23. S. Luti, L. Sella, A. Quarantin, L. Pazzagli, I. Baccelli, Twenty years of research on cerato-platanin family proteins: clues, conclusions, and unsolved issues. Fungal Biology Reviews 34, 13–24 (2020).

24. C. Veneault-Fourrey et al., Genomic and transcriptomic analysis of Laccaria bicolor CAZome reveals insights into polysaccharides remodelling during symbiosis establishment. Fungal Genet. Biol. 72, 168–181 (2014).

25. D. Vincent et al., Secretome of the free-living mycelium from the ectomycorrhizal basidiomycete Laccaria bicolor. Journal of Proteome Research 11, 157–171 (2012).

26. M. Saloheimo et al., Swollenin, a Trichoderma reesei protein with sequence similarity to the plant expansins, exhibits disruption activity on cellulosic materials. Eur. J. Biochem. 269, 4202–4211 (2002).

27. Y. Brotman, E. Briff, A. Viterbo, I. Chet, Role of swollenin, an expansin-like protein from Trichoderma, in plant root colonization. Plant Physiol. 147, 779–789 (2008).

28. V. S. Bharadwaj, B. C. Knott, J. Ståhlberg, G. T. Beckham, M. F. Crowley, The hydrolysis mechanism of a GH45 cellulase and its potential relation to lytic transglycosylase and expansin function. J. Biol. Chem. 295, 4477–4487 (2020).

29. E. Vlasenko, M. Schülein, J. Cherry, F. Xu, Substrate specificity of family 5, 6, 7, 9, 12, and 45 endoglucanases. Bioresour. Technol. 101, 2405–2411 (2010).

30. T. Nomura et al., High-resolution crystal structures of the glycoside hydrolase family 45 endoglucanase EG27II from the snail Ampullaria crossean. Acta Crystallographica. Section D, Structural Biology 75, 426–436 (2019).

31. A. S. Godoy et al., Structure, computational and biochemical analysis of PcCel45A endoglucanase from Phanerochaete chrysosporium and catalytic mechanisms of GH45 subfamily C members. Scientific Reports 8, 3678 (2018).

32. G. J. Davies, S. P. Tolley, B. Henrissat, C. Hjort, M. Schülein, Structures of oligosaccharide-bound forms of the endoglucanase V from Humicola insolens at 1.9 Å resolution. Biochemistry 34, 16210–16220 (1995).

33. A. Nakamura et al., “Newton’s cradle” proton relay with amide-imidic acid tautomerization in inverting cellulase visualized by neutron crystallography. Science Advances 1, e1500263 (2015).

34. L. Okmane et al., Glucomannan and beta-glucan degradation by Mytilus edulis Cel45A: Crystal structure and activity comparison with GH45 subfamily A, B and C. Carbohydr. Polym. 277, 118771 (2022).

35. C. Lohoff, P. C. F. Buchholz, M. Le Roes-Hill, J. Pleiss, Expansin engineering database: a navigation and classification tool for expansins and homologues. Proteins 89, 149–162 (2021).

36. S. McQueen-Mason, D. J. Cosgrove, Disruption of hydrogen bonding between plant cell wall polymers by proteins that induce wall extension. Proceedings of the National Academy of Sciences 91, 6574–6578 (1994).

37. T. Imai et al., Disturbance of the hydrogen bonding in cellulose by bacterial expansin. Cellulose 30, 8423–8438 (2023).

38. E. S. Kim, H. J. Lee, W. G. Bang, I. G. Choi, K. H. Kim, Functional characterization of a bacterial expansin from Bacillus subtilis for enhanced enzymatic hydrolysis of cellulose. Biotechnol. Bioeng. 102, 1342–1353 (2009).

39. H. J. Lee, S. Lee, H. J. Ko, K. H. Kim, I. G. Choi, An expansin-like protein from Hahella chejuensis binds cellulose and enhances cellulase activity. Molecules and Cells 29, 379–385 (2010).

40. Y. Duan et al., Real-time adsorption and action of expansin on cellulose. Biotechnology for Biofuels 11, 317 (2018).

41. T. Wang et al., Sensitivity-enhanced solid-state NMR detection of expansin’s target in plant cell walls. Proceedings of the National Academy of Sciences 110, 16444–16449 (2013).

42. Y. B. Park, D. J. Cosgrove, A revised architecture of primary cell walls based on biomechanical changes induced by substrate-specific endoglucanases. Plant Physiol. 158, 1933–1943 (2012).

43. Y. B. Park, D. J. Cosgrove, Changes in cell wall biomechanical properties in the xyloglucan-deficient xxt1/xxt2 mutant of Arabidopsis. Plant Physiol. 158, 465–475 (2012).

44. I. J. Kim et al., Binding characteristics of a bacterial expansin (BsEXLX1) for various types of pretreated lignocellulose. Appl. Microbiol. Biotechnol. 97, 5381–5388 (2013).

45. B. Bunterngsook, L. Eurwilaichitr, A. Thamchaipenet, V. Champreda, Binding characteristics and synergistic effects of bacterial expansins on cellulosic and hemicellulosic substrates. Bioresource Technolgy 176, 129–135 (2015).

46. M. Olarte-Lozano et al., PcExl1 a novel acid expansin-like protein from the plant pathogen Pectobacterium carotovorum, binds cell walls differently to BsEXLX1. PLoS One 9, e95638 (2014).

47. M. Haddad Momeni et al., Insights into the action of phylogenetically diverse microbial expansins on the structure of cellulose microfibrils. Biotechnology for Biofuels and Bioproduts 17, 56 (2024).

48. N. H. Yennawar, L. C. Li, D. M. Dudzinski, A. Tabuchi, D. J. Cosgrove, Crystal structure and activities of EXPB1 (Zea m 1), a beta-expansin and group-1 pollen allergen from maize. Proceedings of the National Academy of Sciences 103, 14664–14671 (2006).

49. T. A. Gorshkova et al., Alpha- and beta-expansins expressed in different zones of the growing root of maize. Russian Journal of Plant Physiology 71, 53 (2024).

50. M. Monschein et al., Loosenin-like proteins from Phanerochaete carnosa impact both cellulose and chitin fiber networks. Appl. Environ. Microbiol. 89, e01863–01822 (2023).

51. D. Floudas et al., The paleozoic origin of enzymatic lignin decomposition reconstructed from 31 fungal genomes. Science 336, 1715–1719 (2012).

52. Y. Fukasawa, Ecological impacts of fungal wood decay types: a review of current knowledge and future research directions. Ecol. Res. 36, 910–931 (2021).

53. J. Zhang, A. T. Silverstein Kevin, D. Castaño Jesus, M. Figueroa, J. S. Schilling, Gene regulation shifts shed light on fungal adaption in plant biomass decomposers. mBio 10, 10.1128/mbio.02176-02119 (2019).

54. K. Umezawa, M. Niikura, Y. Kojima, B. Goodell, M. Yoshida, Transcriptome analysis of the brown rot fungus Gloeophyllum trabeum during lignocellulose degradation. PLOS ONE 15, e0243984 (2020).

55. G. N. Presley, J. S. Schilling, Distinct growth and secretome strategies for two taxonomically divergent brown rot fungi. Appl. Environ. Microbiol. 83, e02987–02916 (2017).

56. G. N. Presley, E. Panisko, S. O. Purvine, J. S. Schilling, Coupling secretomics with enzyme activities to compare the temporal processes of wood metabolism among white and brown rot fungi. Appl. Environ. Microbiol. 84, e00159–00118 (2018).

57. J. Abramson et al., Accurate structure prediction of biomolecular interactions with AlphaFold 3. Nature 630, 493–500 (2024).

58. S. Ding et al., Boosting enzymatic degradation of cellulose using a fungal expansin: structural insight into the pretreatment mechanism. Bioresour. Technol. 358, 127434 (2022).

59. D. Dahiya et al., SANS investigation of fungal loosenins reveals substrate-dependent impacts of protein action on the inter-microfibril arrangement of cellulosic substrates. Biotechnology for Biofuels and Bioproducts 18, 27 (2025).

60. R. M. Castillo et al., A six-stranded double-psi beta-barrel is shared by several protein superfamilies. Structure 7, 227–236 (1999).

61. CAZypedia (2021) Glycoside Hydrolase Family 45 --- CAZypedia{,} © 2007-2021 The Authors and Curators of CAZypedia.

62. O. A. Hegnar et al., Quantifying oxidation of cellulose-associated glucuronoxylan by two lytic polysaccharide monooxygenases from Neurospora crassa. Appl. Environ. Microbiol. 87, e0165221 (2021).

63. M. Frommhagen et al., Discovery of the combined oxidative cleavage of plant xylan and cellulose by a new fungal polysaccharide monooxygenase. Biotechnology for Biofuels and Bioproducts 8, 101 (2015).

64. J. R. Bromley et al., GUX1 and GUX2 glucuronyltransferases decorate distinct domains of glucuronoxylan with different substitution patterns. The Plant Journal 74, 423–434 (2013).

65. Y. Yoshimi, T. Tryfona, P. Dupree, Structure, modification pattern, and conformation of hemicellulose in plant biomass. Journal of Applied Glycoscience 72 (2024).

66. A. Kirui et al., Carbohydrate-aromatic interface and molecular architecture of lignocellulose. Nature Communications 13, 538 (2022).

67. O. M. Terrett et al., Molecular architecture of softwood revealed by solid-state NMR. Nature Communications 10, 4978 (2019).

68. S. Malgas, M. S. Mafa, L. Mkabayi, B. I. Pletschke, A mini review of xylanolytic enzymes with regards to their synergistic interactions during hetero-xylan degradation. World J. Microbiol. Biotechnol. 35, 187 (2019).

69. T. Wang, Y. Chen, A. Tabuchi, D. J. Cosgrove, M. Hong, The target of β-Expansin EXPB1 in maize cell walls from binding and solid-state NMR studies. Plant Physiol. 172, 2107–2119 (2016).

70. K. Gourlay et al., Swollenin aids in the amorphogenesis step during the enzymatic hydrolysis of pretreated biomass. Bioresour. Technol. 142, 498–503 (2013).

71. M. Andberg, M. Penttilä, M. Saloheimo, Swollenin from Trichoderma reesei exhibits hydrolytic activity against cellulosic substrates with features of both endoglucanases and cellobiohydrolases. Bioresour. Technol. 181, 105–113 (2015).

72. D. J. Cosgrove, Z. C. Li, Role of expansin in cell enlargement of oat coleoptiles (analysis of developmental gradients and photocontrol). Plant Physiol. 103, 1321–1328 (1993).

73. D. J. Cosgrove, Plant cell wall extensibility: connecting plant cell growth with cell wall structure, mechanics, and the action of wall-modifying enzymes. J. Exp. Bot. 67, 463–476 (2016).

74. I. Baccelli, S. Luti, R. Bernardi, A. Scala, L. Pazzagli, Cerato-platanin shows expansin-like activity on cellulosic materials. Appl. Microbiol. Biotechnol. 98, 175–184 (2014).

75. G. Jäger et al., How recombinant swollenin from Kluyveromyces lactis affects cellulosic substrates and accelerates their hydrolysis. Biotechnology for Biofuels 4, 33 (2011).

76. N. Georgelis, N. Nikolaidis, D. J. Cosgrove, Biochemical analysis of expansin-like proteins from microbes. Carbohydr. Polym. 100, 17–23 (2014).

77. N. J. Grantham et al., An even pattern of xylan substitution is critical for interaction with cellulose in plant cell walls. Nature Plants 3, 859–865 (2017).

78. T. J. Simmons et al., Folding of xylan onto cellulose fibrils in plant cell walls revealed by solid-state NMR. Nature Communications 7, 13902 (2016).

79. X. Kang et al., Lignin-polysaccharide interactions in plant secondary cell walls revealed by solid-state NMR. Nature Communications 10, 347 (2019).

80. M. Busse-Wicher, N. J. Grantham, J. J. Lyczakowski, N. Nikolovski, P. Dupree, Xylan decoration patterns and the plant secondary cell wall molecular architecture. Biochem. Soc. Trans. 44, 74–78 (2016).

81. T. R. I. Cataldi, C. Campa, G. E. De Benedetto, Carbohydrate analysis by high-performance anion-exchange chromatography with pulsed amperometric detection: the potential is still growing. Fresenius J. Anal. Chem. 368, 739–758 (2000).

