## Supporting Information for "A single-domain expansin-like protein from *Gloeophyllum trabeum* able to cleave xylan"

PNAS

* Anikó Várnai

**This PDF file includes:**

Supporting text
Figures S1 to S26
Tables S1 to S6
SI References

### Supporting Information Text

### Materials and methods

#### Sequence and structure analysis

The sequence of *Gt*EXPN_133317 from *Gloeophyllum trabeum* (GenBank ID, XP_007870440.1; UniProt ID, S7PTB8), together with all EXPN proteins from the genomes of *G. trabeum* and nine selected brown-rot fungal species, was retrieved from the JGI Mycocosm website (https://mycocosm.jgi.doe.gov/mycocosm/home) (see next section). Multiple sequence alignment (MSA) of EXPNs was performed with EMBL-EBI's online MUSCLE tool (https://www.ebi.ac.uk/jdispatcher/msa/muscle). Structure-based MSA of ERPs and GH45 proteins was performed with T-Coffee's Expresso server (1), and the alignment in the IV-V loop (between the β4 and β5 strands) was corrected based on the PDB structures of the aligned proteins. The AlphaFold model of the mature *Gt*EXPN_133317 protein was predicted using AlphaFold 3 (https://alphafoldserver.com/). Structure analyses and visualization were performed using PyMol (The PyMOL Molecular Graphics System, Version 2.4.1, Schrödinger, LLC). Molecular docking simulations were performed with SwissDock and AutoDock Vina (https://www.swissdock.ch/). The Expasy ProtParam tool (https://web.expasy.org/protparam/) was used to determine the theoretical pI and predicted molecular mass of *Gt*EXPN_133317. The NetNGlyc-1.0 (https://services.healthtech.dtu.dk/services/NetNGlyc-1.0/) and NetOGlyc-4.0 (https://services.healthtech.dtu.dk/services/NetOGlyc-4.0/) servers were used to predict N-glycosylation and O-glycosylation sites, respectively, in *Gt*EXPN_133317.

#### Multiple sequence alignment (MSA) and phylogeny of EXPNs

In order to look for conserved residues among EXPNs in brown-rot fungi, the sequences of EXPNs were collected from the annotated genomes of nine fungal brown-rot strains: *Antrodia sinuosa* LB1 v1.0, *Daedalea quercina* v1.0, *Gloeophyllum trabeum* v1.0, *Laetiporus sulphureus* var. *sulphureus* v1.0, *Postia* (*Rhodonia*) *placenta* MAD 698-R v1.0, *Postia placenta* MAD-698-R-SB12 v1.0, *Serpula lacrymans* S7.3 v2.0, *Serpula lacrymans* S7.9 v2.0, and *Wolfiporia cocos* MD-104 SS10 v1.0. The DPBB domains (InterPro, IPR036908) of the obtained 125 sequences (without their signal peptide or additional domains) were aligned using EMBL-EBI's online MUSCLE5 tool to generate the phylogenetic tree shown in **Fig. S17**. For reference, we included six DPBB domains with known structure, which are shown in **Fig. 2** (three from plants, one from a bacterium, and two from fungi). In a parallel approach, aimed at the identification of conserved residues in a subset of EXPNs including *Gt*EXPN_133317, the protein sequences were filtered using the following criteria: 1) we omitted fragments of DPBB domains and multidomain proteins (as they formed clearly separate clusters from the one containing *Gt*EXPN_133317); 2) we omitted sequences with 3 or 5 cysteines (NB: The DPBB domains of EXPNs in brown-rot fungi possess two highly conserved disulfide bonds and thus 4 cysteines); 3) we selected the sequences where the four conserved cysteines were located at the same position as in *Gt*EXPN_133317; 4) to further reduce the number of sequences, we selected the sequences that share at least 50% sequence identity with *Gt*EXPN_133317. The DPBB domains thus selected, 24 in total, including two from *G. trabeum*, were then aligned using EMBL-EBI's online MUSCLE tool. The resulting MSA is shown in **Fig. S5**.

#### Protein production in *Pichia pastoris*

The gene encoding *Gt*EXPN_133317 (MycoCosm protein ID, 133317; UniProt ID, S7PTB8) from *G. trabeum* v1.0 (Glotr1.1), including its native signal peptide, was codon optimized for expression in *Pichia pastoris,* and de novo synthesized by GenScript (Piscataway, NJ, USA) into a pPICZ A plasmid. Furthermore, genes encoding *Gt*EXPN_133317 mutants (four in total) were synthesized by GenScript in pPICZ A plasmids. The pPICZ A plasmids containing the inserts were transformed into chemically competent *Escherichia coli* TOP10 cells for plasmid maintenance and production. Recombinant *E. coli* cells were grown overnight on LB broth supplemented with 25 µg/mL zeocin, and plasmids were isolated from the cells using the Wizard® Plus SV Minipreps DNA purification system (Promega, Madison, WI, USA). The isolated plasmids were linearized with *Sac*I restriction enzyme (New England Biolabs, Ipswich, MA, USA) and transformed into *P. pastoris* KM71H (for WT and D87A) or *P. pastoris* GS115 (for D25N, D76N, and D76A) using electroporation according to the manufacturer's recommendations (Invitrogen, Thermo Fisher Scientific, San Diego, CA, USA). Transformants with the highest protein production levels were selected, and the inserted genes were verified by sequencing (Eurofins Genomics).

For protein production, 10 mL Buffered Glycerol-complex Medium (BMGY) was inoculated with the expression strains and incubated overnight at 29 °C in a shaking incubator at 220 rpm (New Brunswick Scientific, NJ, USA). These pre-cultures were used to inoculate 1 L BMGY, and the main cultures were grown at 29 °C in 3 L flasks in a shaking incubator at 220 rpm for 20-22 h, to an $OD_{600}$ of 3-6. The cells were recovered after centrifuging the cultures at 1500 *g* for 5 min at room temperature. KM71H cells were resuspended in 200 mL Buffered Methanol-complex Medium (BMMY) (final $OD_{600} \approx 30$), while GS115 cells were resuspended in 3 L of BMMY (final $OD_{600} \approx 1$), and both were incubated in 2 L baffled flasks covered with 2 layers of sterile gauze at 29 °C in a shaking incubator at 210 rpm, supplementing the cultures with methanol to a final concentration of 0.5% (v/v) every 24 h. After 48 h, the culture broth was centrifuged for 10 min at 3000 *g* and 4 °C, and the supernatant was filtered through a 0.22 µm sterile filter (Merck, Darmstadt, Germany). Filtered supernatants were stored at 4 °C.

**Protein production in *Escherichia coli***

The gene sequence encoding the mature *Gt*EXPN_133317 protein was codon optimized for expression in *E. coli* and de novo synthesized, fusing it to the pelB signal peptide, and inserted into the pET-22b (+) plasmid, by GenScript. Nucleotides coding for two extra amino acids (M and D) were incorporated between the signal peptide-encoding DNA sequence and the gene. The plasmid was transformed into chemically competent *E. coli* One Shot™ BL21(DE3) cells (Invitrogen) for expression, and the inserted genes were verified by sequencing (Eurofins Genomics).

Protein production was performed as described by Courtade et al. (2), with the following modifications. Pre-cultures (60 mL LB broth supplemented with 100 µg/mL ampicillin in 150 mL baffled flasks) were inoculated with recombinant cells and incubated overnight at 37 °C in a shaking incubator at 250 rpm. The pre-cultures were transferred into 1 L LB broth supplemented with 100 µg/mL ampicillin in 3 L baffled flasks, and the main cultures were incubated further at 37 °C in a shaking incubator at 250 rpm until an $OD_{600}$ of 0.4. Protein production was then induced by adding isopropyl β-D-1-thiogalactopyranoside (IPTG) at a final concentration of 0.5 µM, followed by further cultivation at 37 °C in a shaking incubator (250 rpm). At 3 h post-induction, cells were harvested by centrifugation at 8000 *g* and 4 °C for 10 min and subjected to osmotic shock using spheroplast buffer (3) to obtain a periplasmic extract, which was then filtered using a 0.22 µm sterile filter and stored at 4 °C.

**Protein purification**

Filtered *P. pastoris* culture broths and *E. coli* periplasmic extracts were concentrated approximately 12 and 6-fold, respectively, to 100 mL. The concentrated broths and periplasmic extracts were then washed three times with 250 mL Milli-Q water and buffer-exchanged into 10 mM Tris/HCl buffer, pH 7.5. Using a Bio-Rad purifier system (Bio-Rad, Hercules, CA, USA), the culture broths or periplasmic extracts were loaded onto a 5-mL HiTrap™ Q Sepharose Fast Flow (FF) column (GE Healthcare, Uppsala, Sweden) equilibrated with 10 mM Tris-HCl buffer, pH 7.5, followed by elution with a linear 0–1 M NaCl gradient at a flow rate of 3 mL/min over 16 column volumes. The fractions containing the protein were identified by SDS-PAGE analysis, after which these were pooled, concentrated to 2 mL, and buffer-exchanged to 20 mM sodium acetate (pH 5.0) or 50 mM BisTris/HCl (pH 6.5) buffer containing 200 mM NaCl, using a 3-kDa Vivaspin centrifugal tube (Merck) at 4 °C and 4500 *g*. The concentrated protein solutions were submitted to size exclusion chromatography using a HiLoad 16/600 Superdex G-75 pg column (Protein Ark) equilibrated with 20 mM sodium acetate (pH 5.0) or 50 mM BisTris/HCl (pH 6.5) buffer with 200 mM NaCl, using a flow rate of 1 mL/min. Fractions containing the protein were pooled and concentrated to a final volume of 2 mL in 20 mM sodium acetate (pH 5.0), 50 mM BisTris/HCl (pH 6.5), 25 mM MES/KOH (pH 6.5), 25 mM HEPES/KOH (pH 7.5), or 25 mM Tris/HCl (pH 9.5), as described above. Protein purity was assessed by SDS-PAGE (**Figures S7** and **S20**). The concentration of purified proteins was determined with the Bradford assay (4), using bovine serum albumin (BSA) as protein standard. The yields of proteins recovered after purification are shown in **Table S6**. Theoretical protein molecular mass values, calculated using Expasy's ProtParam (https://web.expasy.org/protparam/) tool and taken into account the formation of two disulfide bridges in the mature protein, were as follows: *Gt*EXPN_133317$^{Ec}$ wildtype, 12707.99 Da; *Gt*EXPN_133317$^{Pp}$ wildtype, 12461.71 Da; *Gt*EXPN_133317$^{Pp}$ D25N, 12460.72 Da; *Gt*EXPN_133317$^{Pp}$ D87A, 12417.70 Da. Note that the *E. coli*-produced protein has a higher molecular mass because of the insertion of an additional Met-Asp after the pelB leader sequence, which is included at the N terminus of the mature protein.

**Materials**

The following substrates were used in this work: phosphoric acid-swollen cellulose (PASC) prepared from Avicel as described by Wood et al. (5), Avicel microcrystalline cellulose (50 µm), carboxymethylcellulose (CMC), Whatman No. 1 filter paper (0.5 mm discs), xyloglucan (XG) from tamarind seed, medium-viscosity mixed-linkage β-(1→3),(1→4)-glucan from barley (MLG), polygalacturonic acid (PGA) from citrus pectin, mannan from ivory nut, (deacetylated) glucuronoxylan from beechwood (with 4-*O*-methylglucuronyl substitutions and without acetylations), xylopentaose and xylohexaose. Standards included glucose, cellobiose, cellotriose, cellotetraose, cellopentaose, cellohexaose, xylose, xylobiose, xylotriose, xylotetraose, xylopentaose, and xylohexaose. All of the substrates and standards except Avicel (Sigma Aldrich) were purchased from Megazyme (Wicklow, Ireland).

**Thermal stability**

Thermal stability was assessed using a thermal shift assay (6). The reaction was composed of 50 µL pre-mix of 60 µM protein, 12.5 µL Milli-Q water, 25 µL of buffer and 12.5 µL 8X SYPRO Orange protein fluorescent dye (Thermo Fisher Scientific) in 20 mM sodium acetate, pH 5.0. Samples were prepared in quadruplicates on a 96-well plate, keeping the plate on ice and in darkness all the time. The plate was sealed with a transparent film and centrifuged for 1 min at 1000 *g*. The samples were heated from 24 to 98 °C at 1.42 °C /min in a StepOnePlus™ Real-Time PCR System (ThermoScientific), and fluorescent scans were recorded every 17 seconds. The apparent melting temperature ($T_m$) was obtained from the derivative curve of each sample.

**Activity assays**

Proteins (1, 10, or 50 µM) were incubated with substrate (2 mg/mL, unless indicated otherwise) in 20 mM sodium acetate (pH 5.0) or, sometimes, as indicated, 50 mM BisTris/HCl (pH 6.5), 25 mM MES/KOH (pH 6.5), 25 mM HEPES/KOH (pH 7.5), or 25 mM Tris/HCl (pH 9.5) buffer. For reactions containing both PASC and xylan, each was added at 1 mg/mL (0.1%, w/v). Reactions with 1 mL total volume were set up in duplicates or triplicates in 15 mL parafilm-sealed glass tubes and were incubated with an inclination of 30° in a shaking incubator at 160 rpm, 200 rpm or 300 rpm and 30 or 37 °C (as indicated) for up to 48 h. If containing both substrates, a pre-incubation before protein addition was performed, at 220 rpm and 37 °C for 20 min. After incubation, samples were filtered, and the supernatants were stored at 4 °C before further analysis. When measuring activity over time (**Figure S21**), enzymes were inactivated after incubation by diluting samples two-fold with 0.2 M NaOH to reach a final NaOH concentration and volume of 100 mM and 100 µL, respectively.

**Detection of reducing sugars**

Total reducing sugars were determined colorimetrically using dinitrosalicylic acid (DNS) (7), using glucose as standard. In brief, 0.36 mL DNS reagent was mixed with 0.36 mL sample solution in a 1.5 mL Eppendorf tube followed by heating at 90 °C for 10 min. After cooling to room temperature, 0.12 mL 40% (w/v) sodium potassium tartrate solution was added to the mixture and absorbance was determined at 575 nm.

**Rheology**

The rheological behavior of reaction mixtures was determined by continuous rotational rheometry, using an Anton Paar MCR-301 rheometer (Anton Paar GmbH, Graz, Austria), with a semi-conical plate geometry (CP50-1, d= 0.101 mm) and a Peltier element keeping the temperature at 20 °C. Constant shear rates ($\gamma$) of 50 $s^{-1}$ and 100 $s^{-1}$ were applied on a 1 mL reaction mixture for 200, 500 and 1000 s. The shear stress ($\tau$) was measured every 10 s, and the apparent viscosity ($\eta$) was determined as the ratio of $\tau$ and $\gamma$.

**Turbidity assessment**

The turbidity of the reaction mixtures was measured spectrophotometrically at 500 and 600 nm using a Cary 60 UV-Vis Spectrophotometer (Agilent Technologies).

**Chromatographic analysis of product profile by HPAEC-PAD**

Reaction products were analyzed using high-performance anion exchange chromatography with pulsed amperometric detection (HPAEC-PAD) using a Dionex ICS-5000 system (Thermo Fisher Scientific). The system was set up with a 3×250 mm Dionex CarboPac PA-200 analytical column and a 3×50 mm guard column (Thermo Fisher Scientific). In some cases (where indicated), the same analysis was done using a Dionex ICS-6000 system (Thermo Fisher Scientific). For the ICS-5000, the employed eluents were prepared as described by Westereng et al. (8), and the flow rate was 0.5 mL/min. For the ICS-6000, the employed eluent was prepared as described by Østby et al. (9), and the flow rate was 63 µL/min. Samples were diluted two-fold in Milli-Q water to a final volume of 100 µL and analyzed using a 39-min separation gradient (10) with the ICS-5000 or a 50-min separation gradient with the ICS-6000 (9). For the identification of native cellulose and xylan-derived oligosaccharides, standards included cellobiose, cellotriose, cellotetraose, cellopentaose, cellohexaose, xylose, xylobiose, xylotriose, xylotetraose, xylopentaose, and xylohexaose. Chromeleon version 7.2.9 (Thermo Fisher Scientific) was utilized to control the system and to analyze the results.

**Analysis of protein quality and reaction products with MALDI-ToF MS**

Reaction products and purified proteins were analyzed by matrix-assisted laser desorption/ionization time-of-flight mass spectrometry (MALDI-ToF MS) UltrafleXtreme mass spectrometer (Bruker Daltonics GmbH, Bremen, Germany) equipped with a Nitrogen 337 nm laser, as described by Agger et al. (11). Carbohydrate and protein samples were mixed in 1:2 and 1:3 (*v*/*v*) ratio, respectively, with a matrix containing 20 mg/mL 2,5-dihydroxybenzoic acid and 1 mM sodium chloride dissolved in 30% (*v*/*v*) acetonitrile and analyzed as described by Hegnar et al. (10).

**Peptide mapping of *Gt*EXPN_133317$^{Pp}$ using LC-MS$^2$**

The procedure described by Tuveng et al. (12) was followed. Briefly, purified *Gt*EXPN_133317$^{Pp}$ was subjected to gel electrophoresis and stained with Coomassie blue, after which the band corresponding to the protein was excised. De-coloring and cleaning of the gel band were carried out at room temperature, the incubation time being 15 min for each step. Subsequently, reduction and alkylation were carried out at room temperature, the incubation time being 30 min for each step. Trypsin (Sequencing grade, Promega, USA) was added to reach a 1:40 (*w*/*w*) ratio relative to the protein concentration. After incubation at 37 °C overnight, peptides were purified using ZipTip C18 pipette tips (Merck Millipore, Cork, Ireland), dried under vacuum (Concentrator plus, Eppendorf, Denmark) and dissolved in 10 mL 2% (*v*/*v*) acetonitrile, 0.1% (*v*/*v*) trifluoroacetic acid. The peptides were analyzed using a nano HPLC-MS/MS system consisting of a Dionex Ultimate 3000 RSLCnano (Thermo Scientific, Bremen, Germany) connected to a Q-Exactive hybrid quadrupole-orbitrap mass spectrometer (Thermo Scientific) equipped with a nano-electrospray ion source, as described by Tuveng et al. (12).

**A**

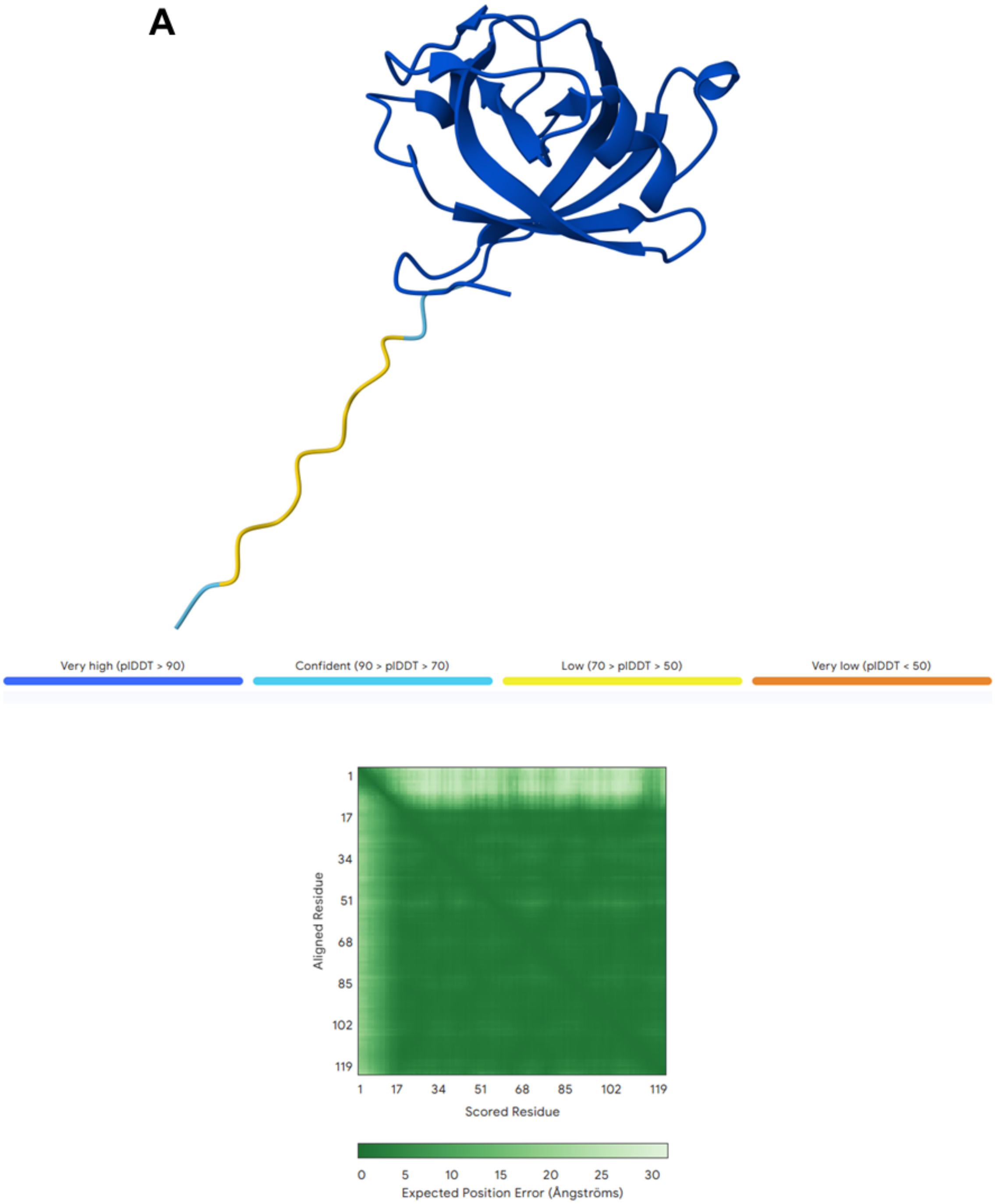

Very high (pIDDT > 90)
Confident (90 > pIDDT > 70)
Low (70 > pIDDT > 50)
Very low (pIDDT < 50)
Aligned Residue
1
17
34
51
68
85
102
119
Scored Residue
0
5
10
15
20
25
30
Expected Position Error (Ångströms)

**B**

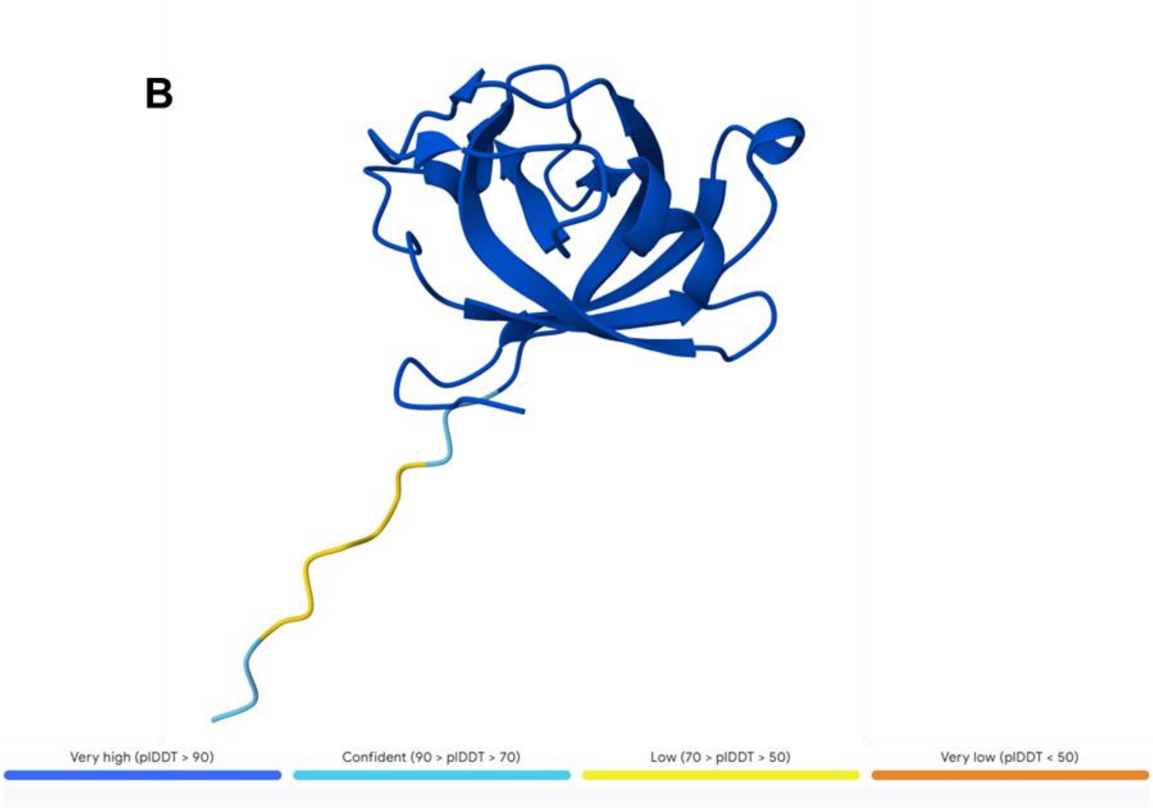

Very high (pIDDT > 90)
Confident (90 > pIDDT > 70)
Low (70 > pIDDT > 50)
Very low (pIDDT < 50)


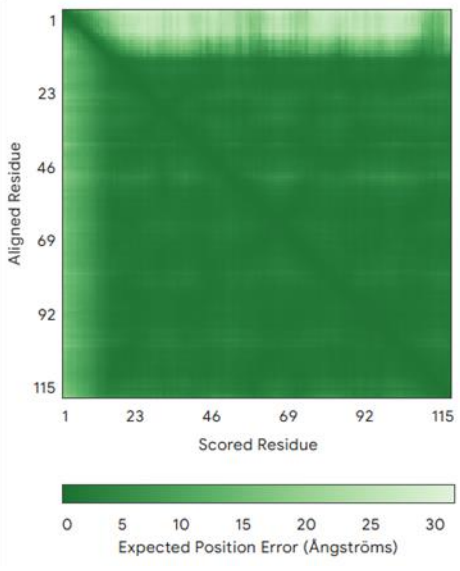

1
23
46
69
92
115
Aligned Residue
Scored Residue
0 5 10 15 20 25 30
Expected Position Error (Ångströms)

**Figure S1. The Alphafold3 predicted structures of A) *Gt*EXPN_133317 WT and B) *Gt*EXPN_133317$^{Pp}$ WT.** Each figure displays: the predicted structural model of the protein with a color-code indicating the Predicted Local Distance Difference Test (pLDDT) scores and a Predicted Aligned Error (PAE) heatmap showing pairwise predicted errors. These analyses document high predicted model accuracies for the complete protein, except for the N-terminal tail that likely is flexible and disordered. The predicted structures of all mutants discussed in this study, including those with mutations at position 76 were essentially identical to the structures shown in the figure and came with similar predicted accuracies. See **Figures S2, S3** and **S4** for a comparison of the active site regions and substrate-binding surfaces of *Gt*EXPN_133317 and resolved protein structures of expansin-related proteins and GH45s.

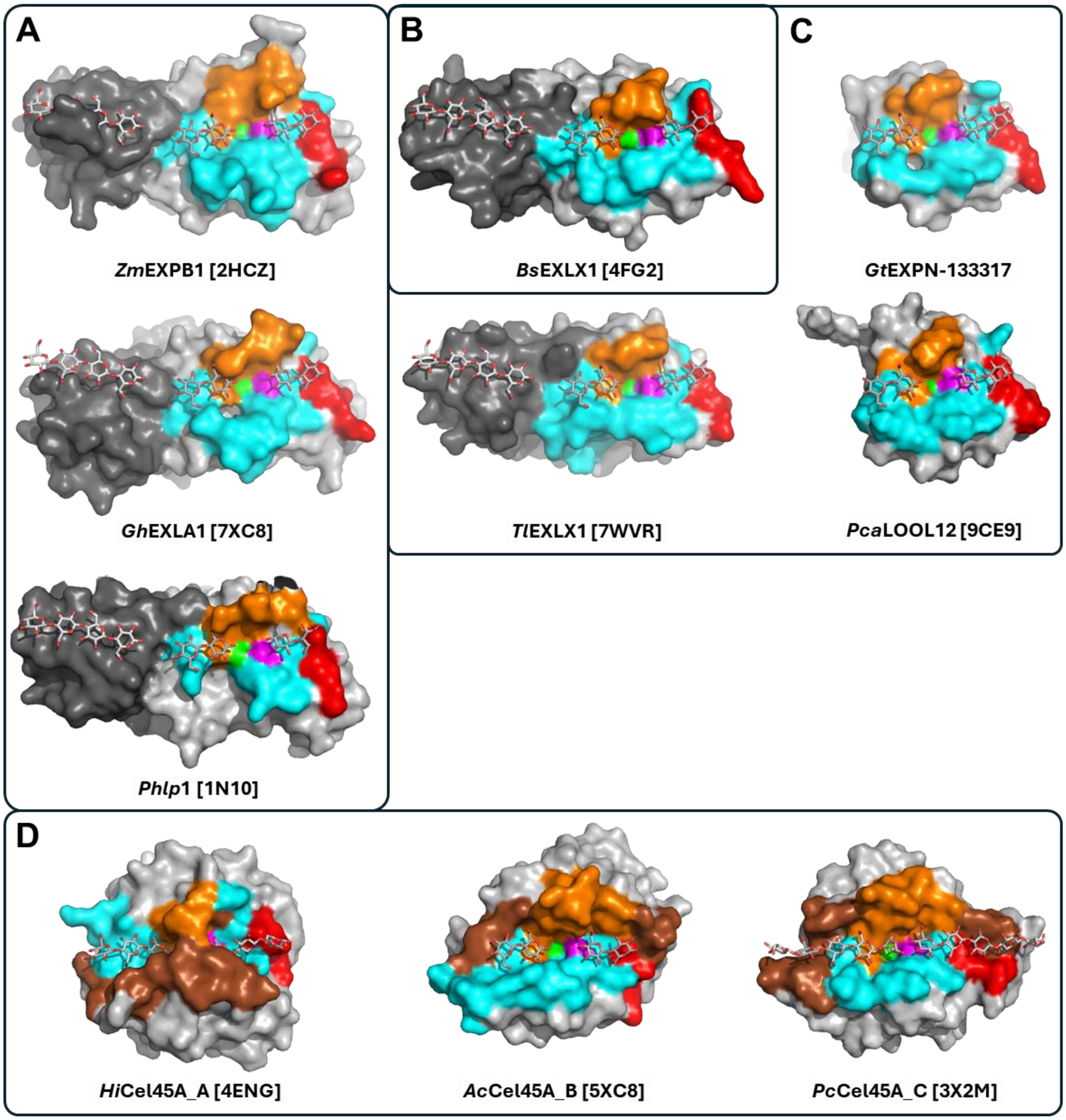


**Figure S2. Active-site segments of plant (A), bacterial (B) and fungal (C) ERPs in comparison with GH45 endoglucanases (D).** The segments were coloured according to the annotation provided in **Figure 2**. The conserved aspartate is shown in magenta; the alanine forming the bottom of a pocket next to the aspartate is shown in green. For comparison purposes, the two cellobiose molecules found in the structure of *Ac*Cel45A [5XC8] are shown at the catalytic site for all ERP structures. Furthermore, a cellotetraose bound to the CBM63 domain in the structure of *Bs*EXLX1 [4FG2] is shown for all plant and bacterial expansins carrying a CBM63. The structures of *Hi*Cel45A [4ENG], *Ac*Cel45A [5XC8], and *Pc*Cel45A [3X2M] are shown with two cellotriose, cellobiose, and cellopentaose molecules, respectively, which are part of the corresponding structures. PDB IDs are indicated in square brackets for all proteins with a known crystal structure. The structure of *Gt*EXPN_133317 was predicted using AlphaFold3 (see **Figure S1** for confidence scores). This figure indicates clear differences in the shape of the substrate-binding surface and, consequently, differences in the proteins' abilities to accommodate linear (and potentially branched) polysaccharides.

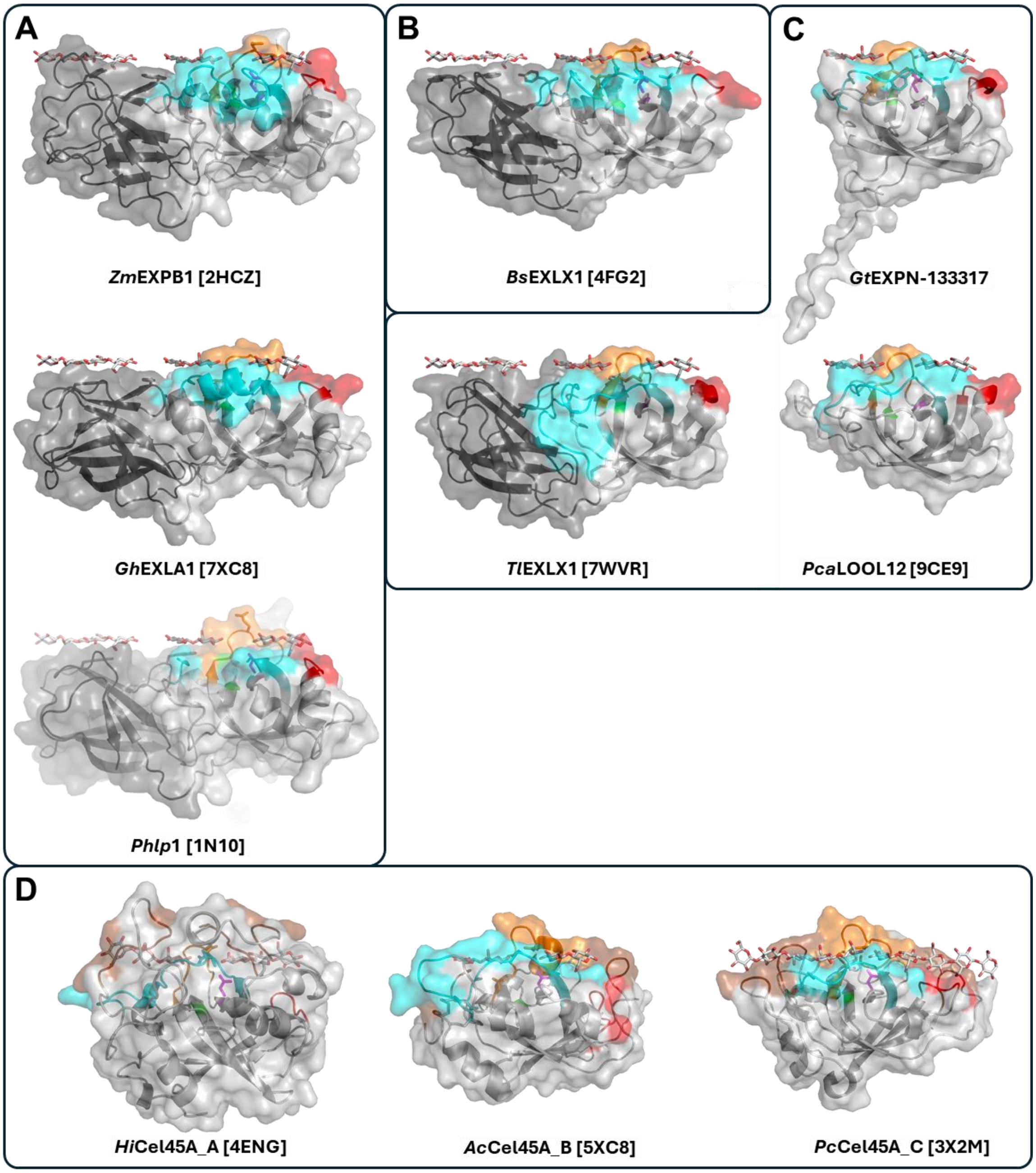


**Figure S3. Sideview of plant (A), bacterial (B) and fungal (C) expansins and comparison with GH45 endoglucanases (D).** These structural models are the same as those shown in **Figure S2**.

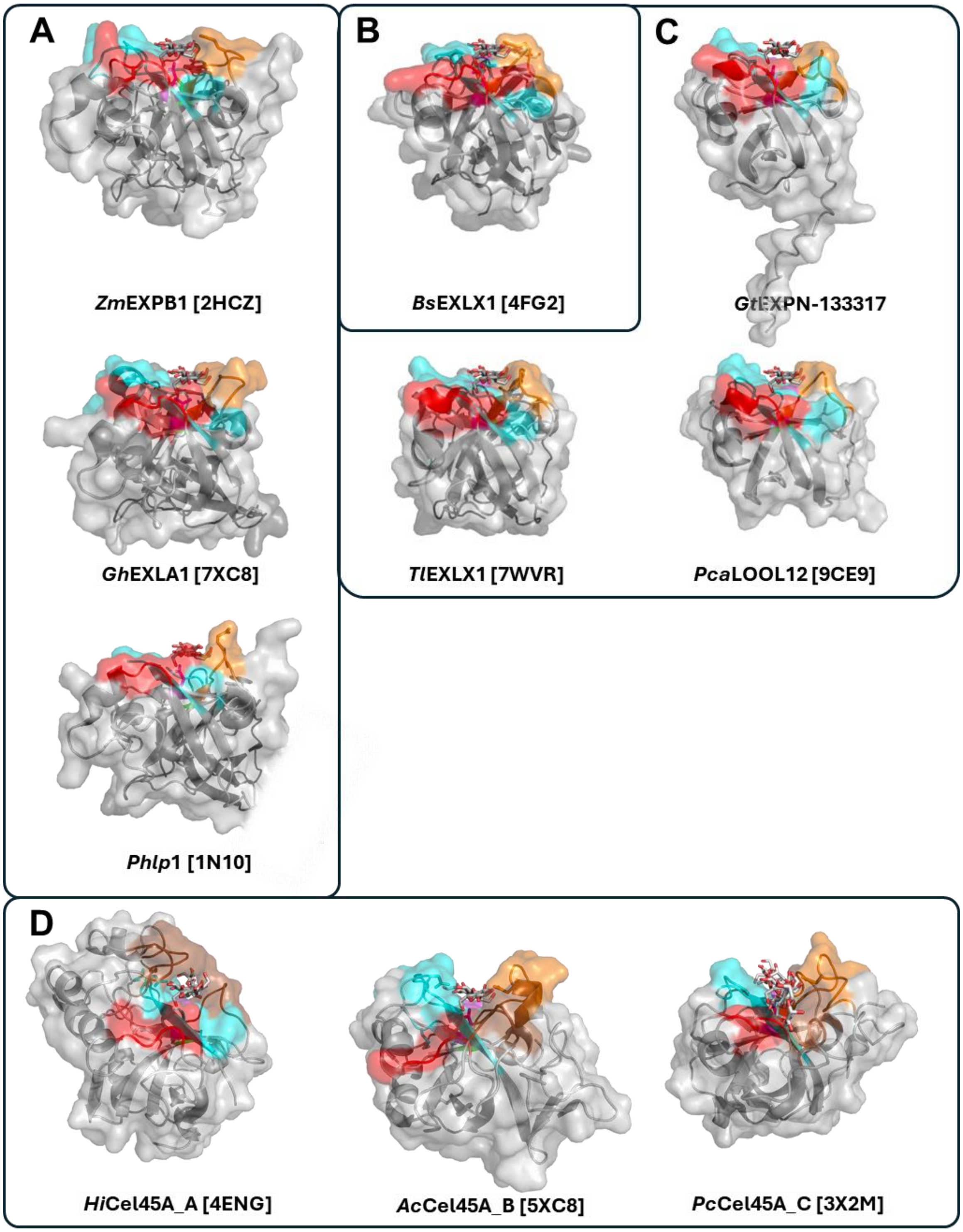


**Figure S4. The DPBB domains of plant (A), bacterial (B) and fungal (C) expansins and of GH45 endoglucanases (D).** See **Figure S2** for details. This figure shows clear differences in the shape of the substrate-binding surface, which likely reflect differences in the proteins' abilities to accommodate linear and branched polysaccharides.

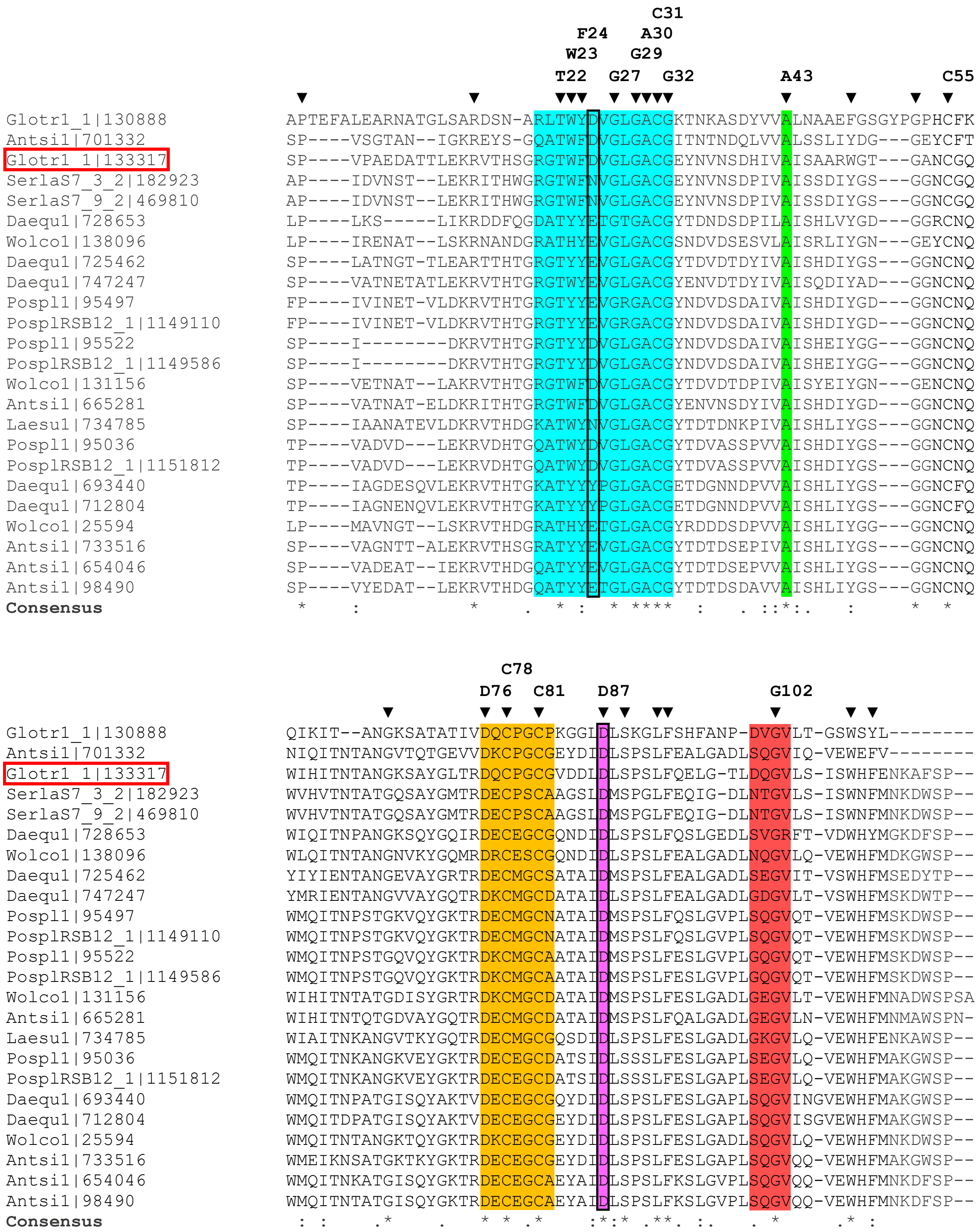


**Figure S5. Multiple sequence alignment of selected brown-rot fungal EXPNs.** *Gt*EXPN_133317 is indicated in a red square. Substrate-binding segments are marked with the background colours as indicated in **Figure 2**. Fully conserved residues and conserved aromatic residues are marked with a black triangle; those potentially involved in substrate binding or in disulfide bridges are indicated above the alignment. A conserved Asp (Asp87) and a conserved Ala (Ala43) appear on a magenta and green background, respectively (i.e. same colouring as in other Figures). The catalytic residues discussed in this study (Asp25 and Asp87) are indicated in black boxes.

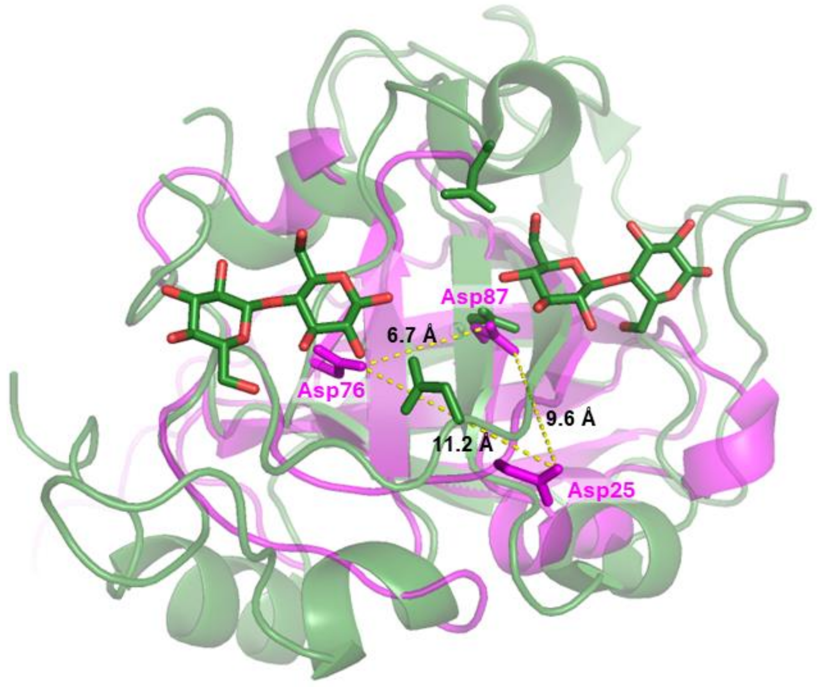


**Figure S6. Distances between three aspartates in *Gt*EXPN_133317.** The conserved aspartates are labelled in the AlphaFold-predicted structure of *Gt*EXPN_133317 (magenta); and the distances between these aspartates are shown in Ångström, as measured using PyMOL. The structure of *Ac*Cel45A in complex with cellobiose [PDB, 5XC8] (green) is superimposed, as shown in **Figure 1**.

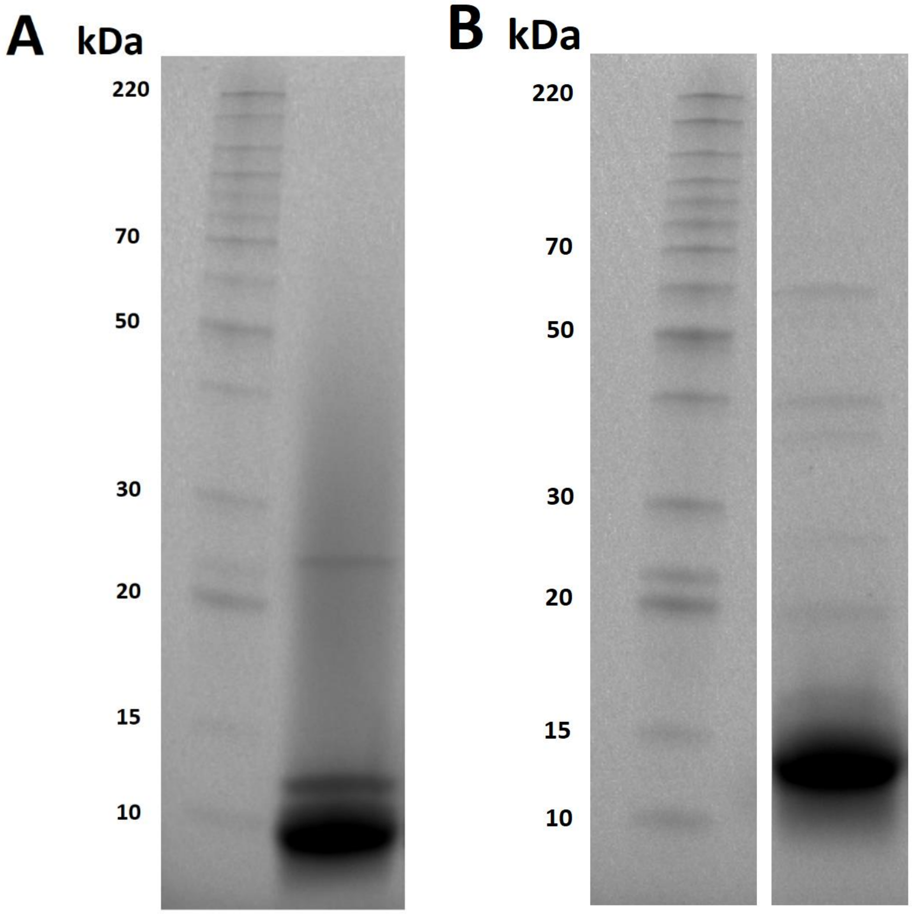


**Figure S7. SDS-PAGE of purified *Gt*EXPN_133317 produced in A) *P. pastoris* and B) *E. coli*.** Each gel shows a lane with BenchMark™ Protein Ladder (left) and a lane with a sample of 40 µg (A) or 50 µg (B) purified *Gt*EXPN_133317 (right). The theoretical molecular masses of the protein variants are (A) 12461.71 Da and (B) 12707.99 Da, as calculated with Expasy's ProtParam tool (https://web.expasy.org/protparam/) and taking into account the formation of two disulfide bridges in the mature protein. The *E. coli*-produced protein has a higher theoretical molecular mass because of the insertion of an additional Met-Asp between the pelB leader sequence and the N-terminus of the mature protein. MALDI-ToF MS analysis of the purified protein preparations showed that the lower apparent molecular mass of the *P. pastoris*-produced protein is due to specific proteolytic processing of the N- and C-termini; see the main text, **Table S4** and **Figures S8 and S9** for details. The molecular masses of marker proteins are indicated on the left side in kilodaltons (kDa). Gel **B)** was cropped to only show relevant gel lanes; both lanes are derived from the same gel and picture.

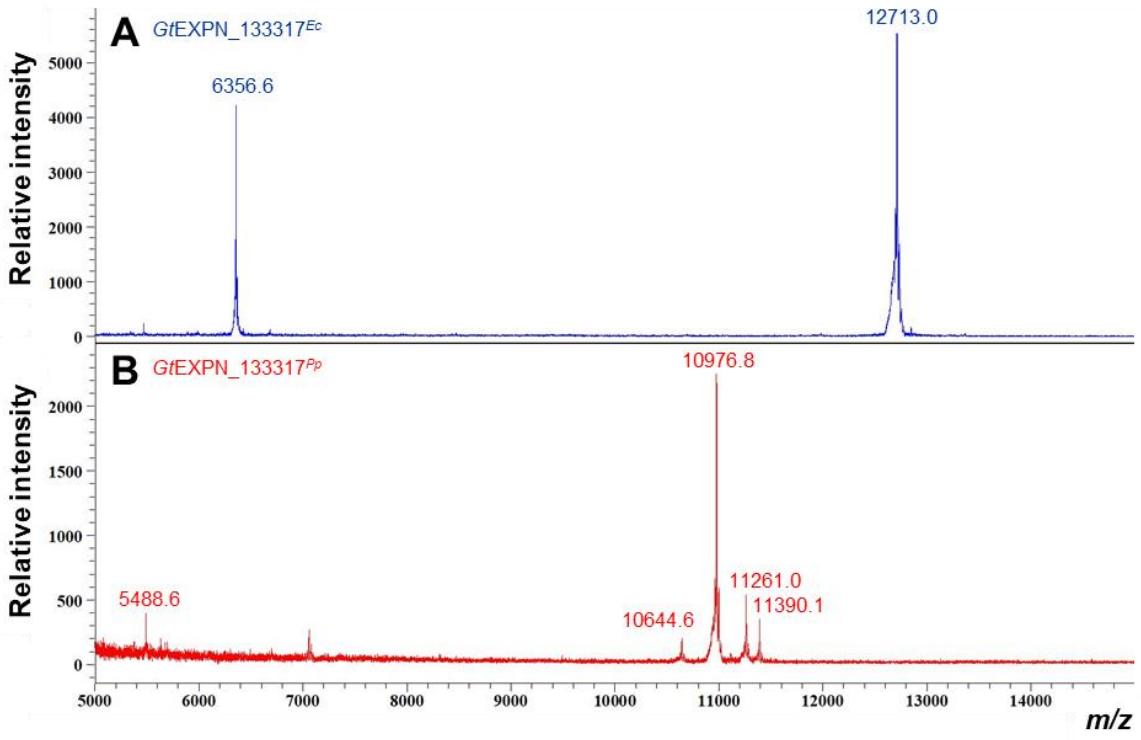


**Figure S8. MALDI-ToF MS analysis of purified A) *Gt*EXPN_133317$^{Ec}$ and B) *Gt*EXPN_133317$^{Pp}$.** For *Gt*EXPN_133317$^{Ec}$, the main *m/z* signals at 12713.0 and 6356.6 correspond to the same protein with a mass of 12713 Da. This value corresponds well to the theoretical molecular mass of *Gt*EXPN_133317$^{Ec}$, which was calculated to be 12707.99 Da using Expasy's ProtParam tool, taking into account the formation of two disulfide bridges in the mature protein (https://web.expasy.org/protparam/). For *Gt*EXPN_133317$^{Pp}$, the main signals at 10976.8 and 5488.6 correspond to the same protein with a mass of 10976.8. Additional peaks indicate the presence of three other proteins with *m/z* values of 10644.6, 11261.0 and 11390.1. The detected apparent masses are smaller than 12461.71 Da, the theoretical molecular mass of *Gt*EXPN_133317$^{Pp}$, as calculated using Expasy's ProtParam tool, taking into account disulfide bridge formation. Detailed analysis of the masses derived from these MS data (see **Table S4**) indicated that these four masses correspond to protein variants of *Gt*EXPN_133317$^{Pp}$ where the N-terminal flexible region (Ser1-Arg14) and/or the short C-terminal tail has been cleaved proteolytically at various positions (**Table S4**, **Figure S9)**. The masses determined here correspond to those observed in the SDS-PAGE analysis shown in **Figure S7**.

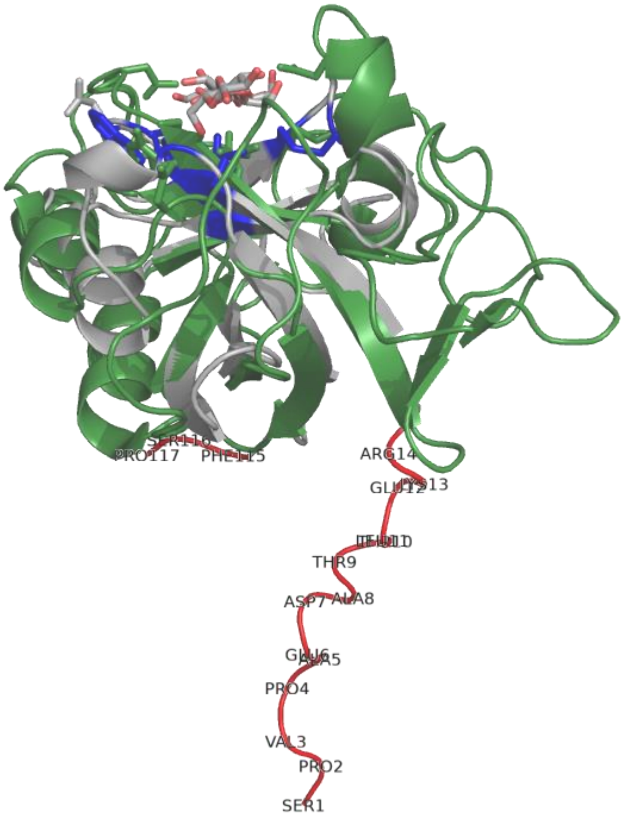


**Figure S9. View of *Gt*EXPN_133317 showing the N- and C-terminal regions that are prone to proteolysis when the protein is produced in *P. pastoris***. The figure shows a structural alignment of *Gt*EXPN_133317 (grey, AlphaFold3 model) and *Ac*Cel45A in complex with cellobiose [5XC8] (green), as in **Figure 1**. The N-terminal (Ser1–Arg14) and C-terminal (Phe115–Pro117) fragments that are proteolytically cleaved during production in *P. pastoris* are shown in red.

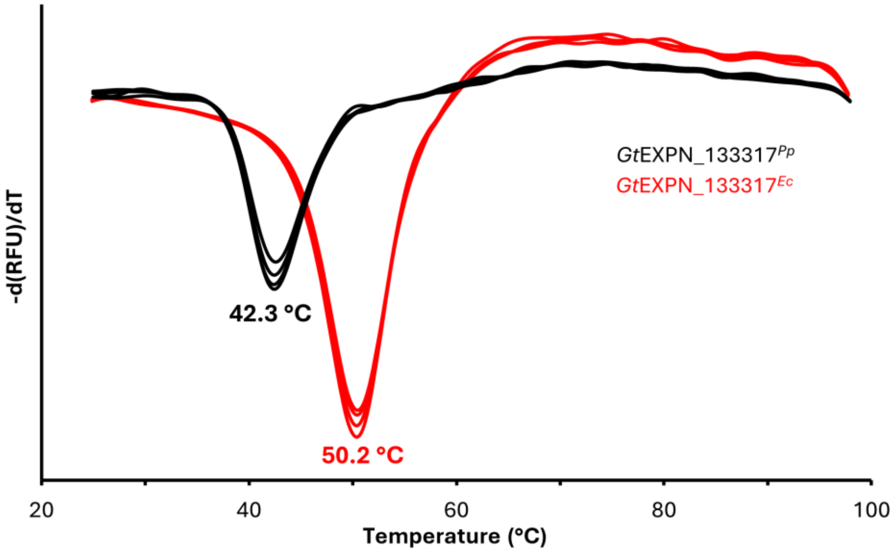


**Figure S10. Thermal stability of *Gt*EXPN_133317.** The figure shows the melting curve and apparent melting temperature ($T_{m\ (app)}$) for *Gt*EXPN_133317$^{Ec}$ (red lines) and *Gt*EXPN_133317$^{Pp}$ (black lines). The derivative of the fluorescence signal ((-d(RFU)/dT), RFU= relative fluorescence units) is plotted against temperature. 30 µM of protein supplemented with SYPRO orange (fluorescent dye) was heated from 24 °C to 98 °C at 1.42 °C/min. Four replicate measurements were performed, which are all shown. The standard deviation in the apparent melting temperature ($T_{m\ (app)}$) was ±0.3 °C. The average $T_{m\ (app)}$ is indicated for each protein. The lower $T_{m\ (app)}$ of *Gt*EXPN_133317$^{Pp}$ indicates reduced stability, likely due to proteolytic cleavage of the *P. pastoris*-produced protein (discussed in the main text and documented in **Table S4** and **Figures S8 & S9**).

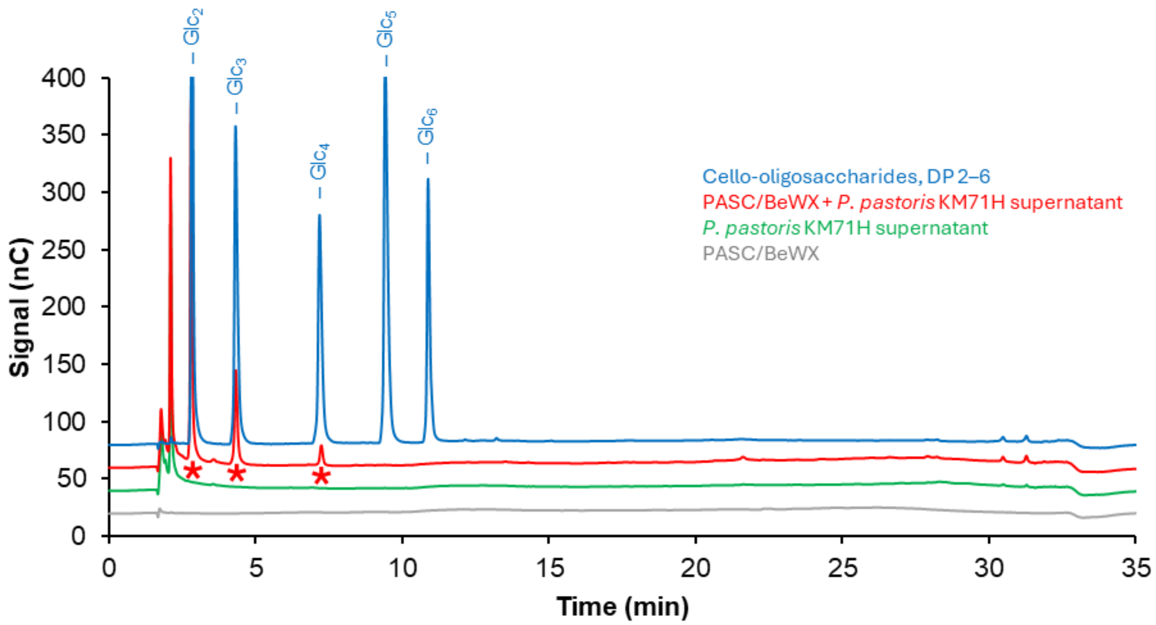


**Figure S11. HPAEC-PAD chromatogram showing endoglucanase background in the concentrated supernatant of *P. pastoris* KM71H.** The red chromatogram shows reaction products after treatment of a mixture of 0.1% (*w*/*v*) beechwood xylan + 0.1% (*w*/*v*) PASC with the 12-fold concentrated supernatant of the *P. pastoris* KM71H strain. Reactions were incubated at 30°C and pH 5.0 for 48 h with horizontal shaking at 300 rpm. Chromatograms for control reactions without substrate (green line) or without *P. pastoris* supernatant (grey line) are also shown. All reactions were performed in triplicates, with similar results; a typical chromatogram is shown. Cello-oligosaccharides with a degree of polymerization (DP) 2–6 (each at 0.05 mg/mL concentration) were used as standard (blue line). The figure shows that *P. pastoris* KM71H exhibits cellulolytic background activity (note the formation of $Glc_2$, $Glc_3$, and $Glc_4$; marked with red asterisks) but not xylanolytic background activity.

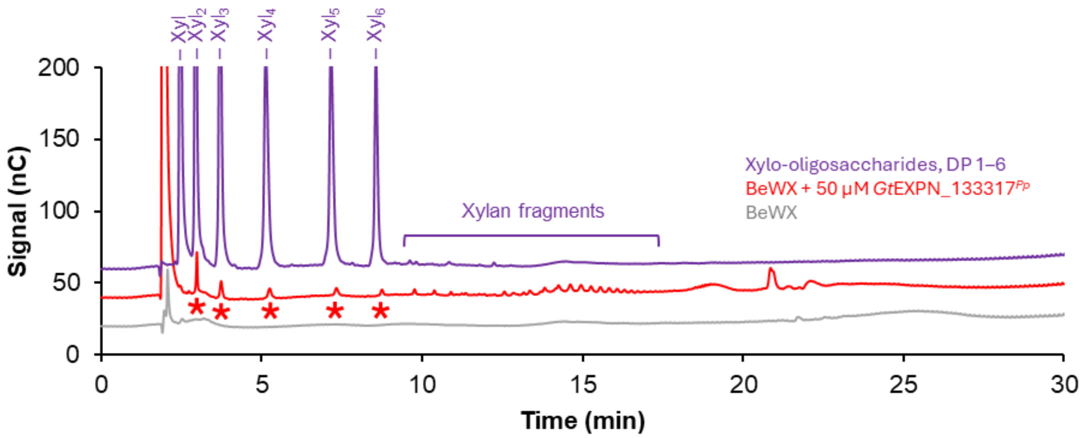


**Figure S12. HPAEC-PAD analysis of products obtained in reactions of *Gt*EXPN_133317*$^{Pp}$* with beechwood xylan only.** In the reactions, 50 µM (red line) *Gt*EXPN_133317*$^{Pp}$* was incubated with 0.2% (*w*/*v*) beechwood xylan at 37°C and pH 5.0 for 48 h with horizontal shaking at 300 rpm. The identified linear xylo-oligosaccharides (with DP 2–6) are marked with red asterisks. Chromatograms for control reactions without enzyme are shown with grey lines. All reactions were performed in triplicates, with similar results; a typical chromatogram is shown. Xylo-oligosaccharides with DP 1–6 (purple), each at 0.025 mg/mL concentration, were used as standard.

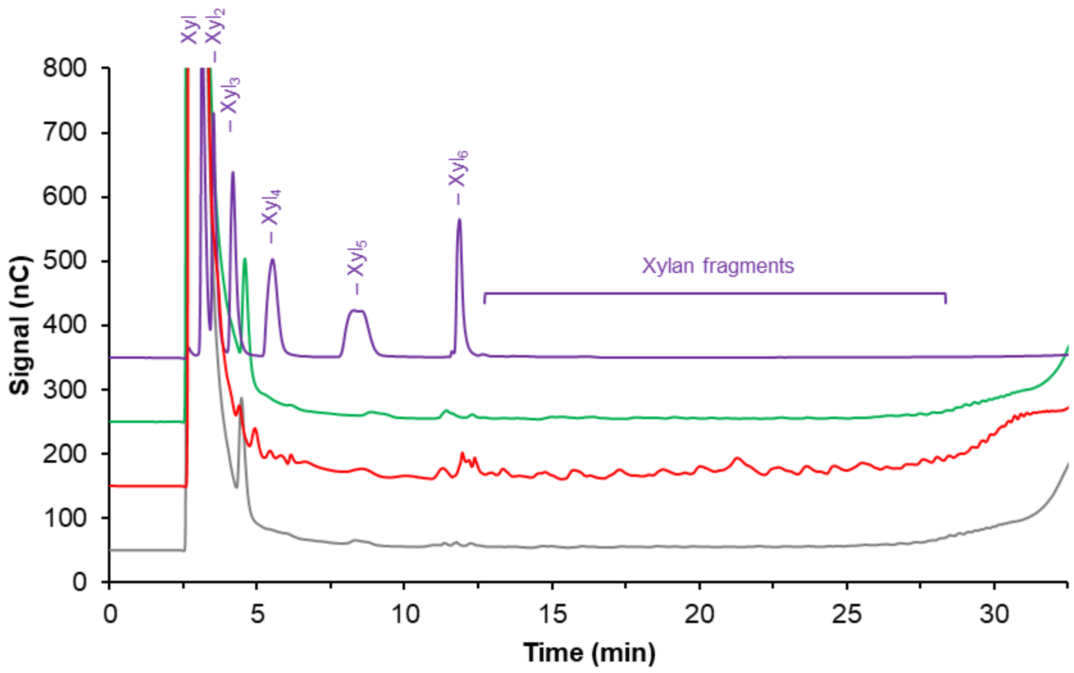


**Figure S13. HPAEC-PAD chromatograms demonstrating inactivation of *Gt*EXPN_133317 after boiling.** In the reactions, 5% (*w*/*v*) beechwood xylan was incubated with 50 µM *Gt*EXPN_133317$^{Ec}$ (red line) or heat-inactivated *Gt*EXPN_133317$^{Ec}$ (green line) at 30 °C, pH 6.5, for 48 h with horizontal shaking at 160 rpm. The result for a control reaction without enzyme is shown with a grey line. All reactions were performed in duplicates, with similar results; a typical chromatogram is shown. Xylo-oligosaccharides with DP 1–6 (purple line), each at 0.025 mg/mL concentration, were used as standard. Reaction products were analyzed using a Dionex ICS-6000 system (as opposed to the ICS-5000 system used in most other experiments) and this leads to changes in retention times.

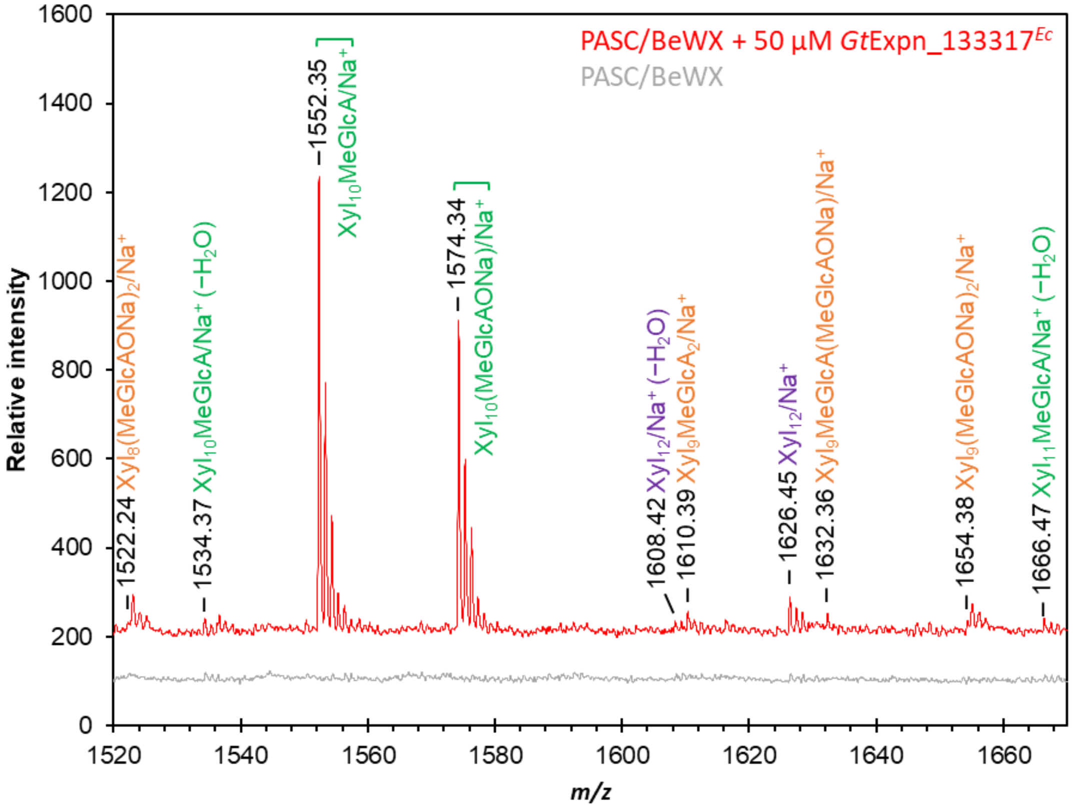


**Figure S14. A close-up of the MALDI-ToF MS spectrum of products generated in the reaction of *Gt*EXPN_133317*Ec* with a 1:1 (*w*/*w*) mixture of PASC and beechwood xylan.** The products include non-substituted (purple labels; $Xyl_n$) and singly (green labels; $Xyl_n$MeGlcA) or doubly (orange labels; $Xyl_n MeGlcA_2$) glucuronylated xylan fragments of varying length, where *n* is the number of xylosyl units in the backbone of the xylan fragments. All species are $Na^+$ adducts ('/$Na^+$') as indicated in the labels. The sodium salt of methyl-glucuronyl side groups is indicated as '(MeGlcAONa)'. Signals for the double $Na^+$-salt of doubly glucuronylated xylo-oligosaccharides [e.g., $Xyl_9(MeGlcAONa)_2/Na^+$] are also visible and labelled. Notably, we also detected species that could correspond to dehydrated species [marked with '($-H_2O$)'] of a number of xylan fragments. The mass spectrum for a control reaction without *Gt*EXPN_133317*Ec* is shown as a grey line. The full spectrum is shown in **Figure 5** of the main manuscript.

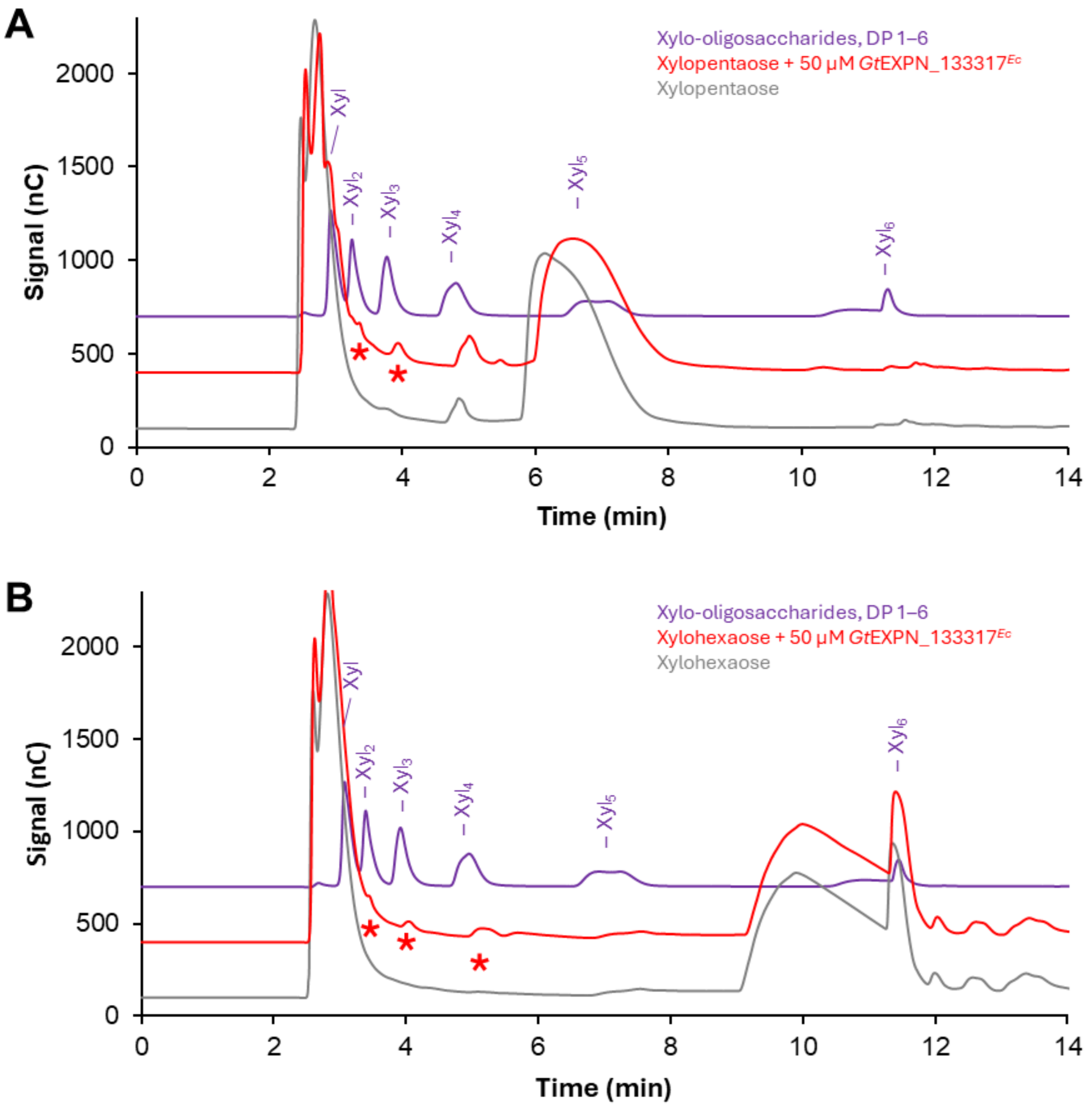


**Figure S15. HPAEC-PAD analysis of products generated in a reaction with *Gt*EXPN_133317$^{Ec}$ and A) xylopentaose and B) xylohexaose.** In the reactions, 50 µM *Gt*EXPN_133317$^{Ec}$ was incubated with 0.1% (*w*/*v*) of xylopentaose (A) or xylohexaose (B) at 30°C, pH 6.5, for 48 h with horizontal shaking at 160 rpm (red chromatograms). Products generated in these reactions are marked with red asterisks. Chromatograms for control reactions without enzyme are shown with grey lines. All reactions were performed in duplicates, with similar results; a typical chromatogram is shown. Xylo-oligosaccharides with DP 1–6 (purple line), each at 0.025 mg/mL concentration, were used as standard. These reaction mixtures, containing high amounts of substrate, were analyzed using the ICS-6000 system, which gave technical challenges in the form of strange peak shapes for the substrate. Despite these issues, the result is clear: small amounts of products are being generated (of note, these are very small amounts relative to the substrate concentration; see main text). MALDI-ToF MS data for these reactions is shown in **Figure S16**.

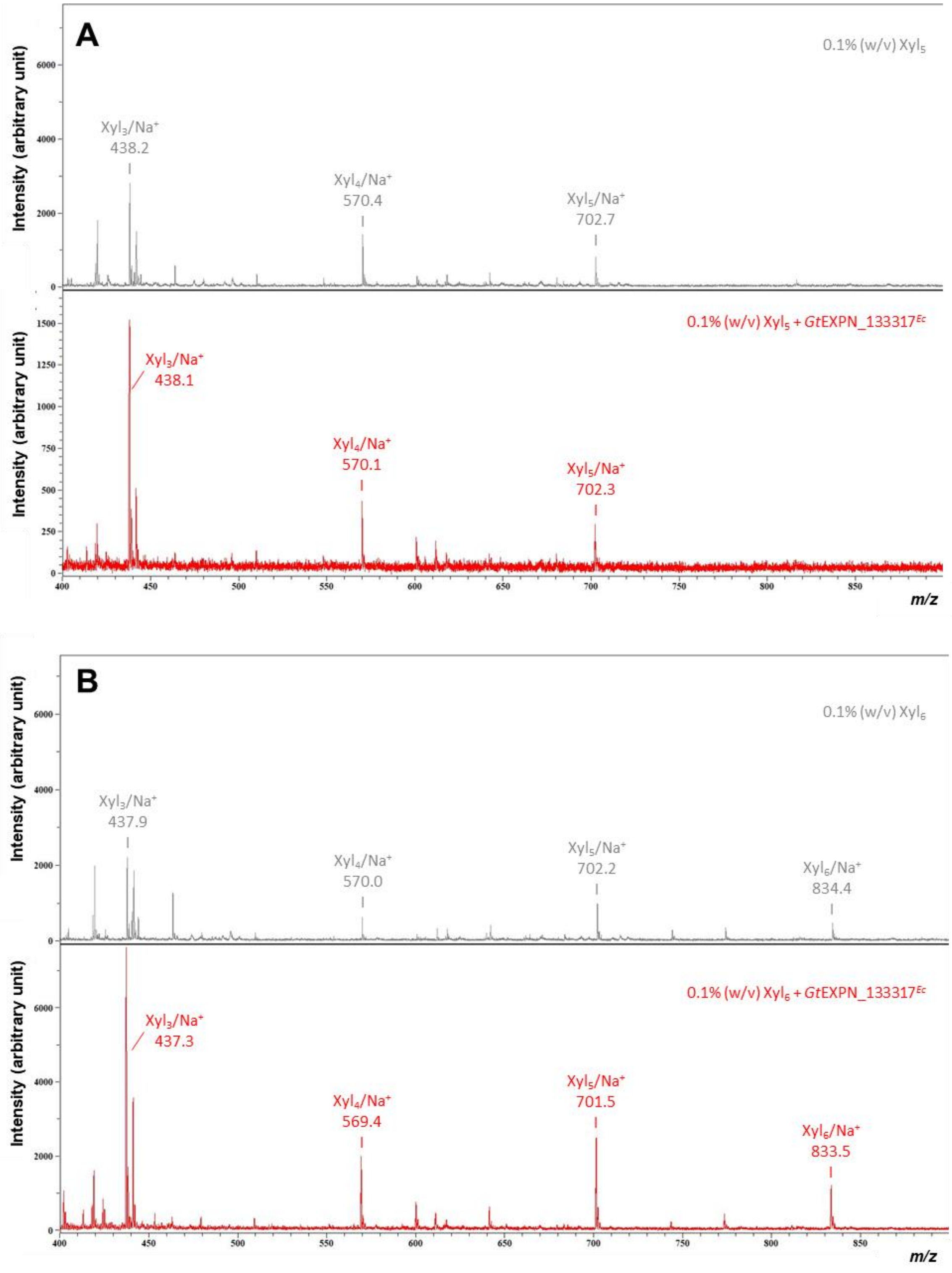


**Figure S16. MALDI-ToF MS analysis of the products generated in a reaction of *Gt*EXPN_133317$^{Ec}$ with xylopentaose (A) and xylohexaose (B).** The reaction conditions are described in the legend of **Fig. S15**. The spectra for control reactions without enzyme is shown in the upper panels (grey spectra). The identified xylo-oligosaccharides (with degree of polymerization, DP, of 3, 4, 5 or 6) are indicated above the signals. The *m/z* values for $Na^+$-adducts are shown. Note that in this type of analysis, the signal intensities do not reflect product abundance, and that quantitative interpretation of these spectra is not possible. For example, the high concentrations of remaining substrate are not reflected in strong *m/z* signals. It is clear, however, that in the reactions with enzyme the signals for the shorter oligomers (= products) increased relative to the signal for the substrate.

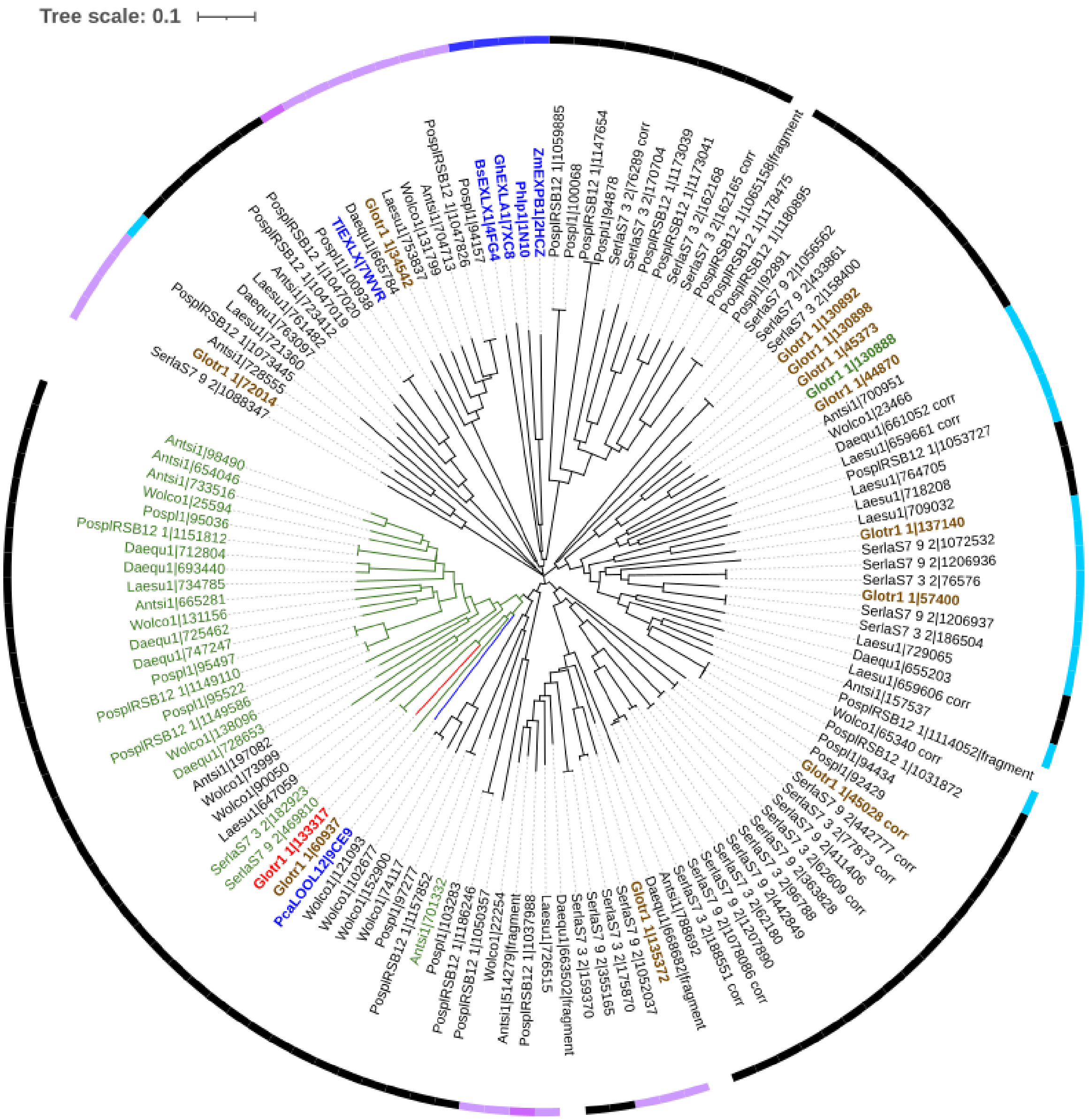


**Figure S17. Phylogenetic tree of the DPBB domains of 125 proteins annotated as EXPN from nine brown-rot fungi.** The sequences were retrieved from JGI's MycoCosm website (https://mycocosm.jgi.doe.gov/fungi/fungi.info.html); signal peptides, predicted with the SignalP-5.0 server (https://services.healthtech.dtu.dk/services/SignalP-5.0/), and additional domains, including the CBM63, were omitted from the sequences. For reference, six DPBB domains of six proteins with resolved crystal structures (*Zm*EXPB1 from *Zea mays* [PDB, 2HCZ], *Gh*EXLA1 from *Gossypium hirsutum* [PDB, 7XC8], *Phl*p1 from *Phleum pratense* [PDB, 1N10], *Bs*EXLX1 from *Bacillus subtilis* [PDB, 4FG4], *Tl*EXLX from *Talaromyces leycettanus* [PDB, 7WVR] and *Pca*LOOL12 from *Phanerochaete carnosa* [PDB, 9CE9]) were included, and their names appear in blue. The sequences selected for multiple sequence alignment (**Figure S5**) based on manual sequence curation are shown in green. EXPNs from *G. trabeum* are shown in brown, except *Gt*EXPN_133317, which is shown in red, and *Gt*EXPN_130888, which is shown in green. In the outer circle, black represents single-domain EXPNs, blue and purple represents multidomain proteins with the DPBB domain at the N- and C-terminus, respectively; dark blue and dark purple indicates the presence of CBM63.

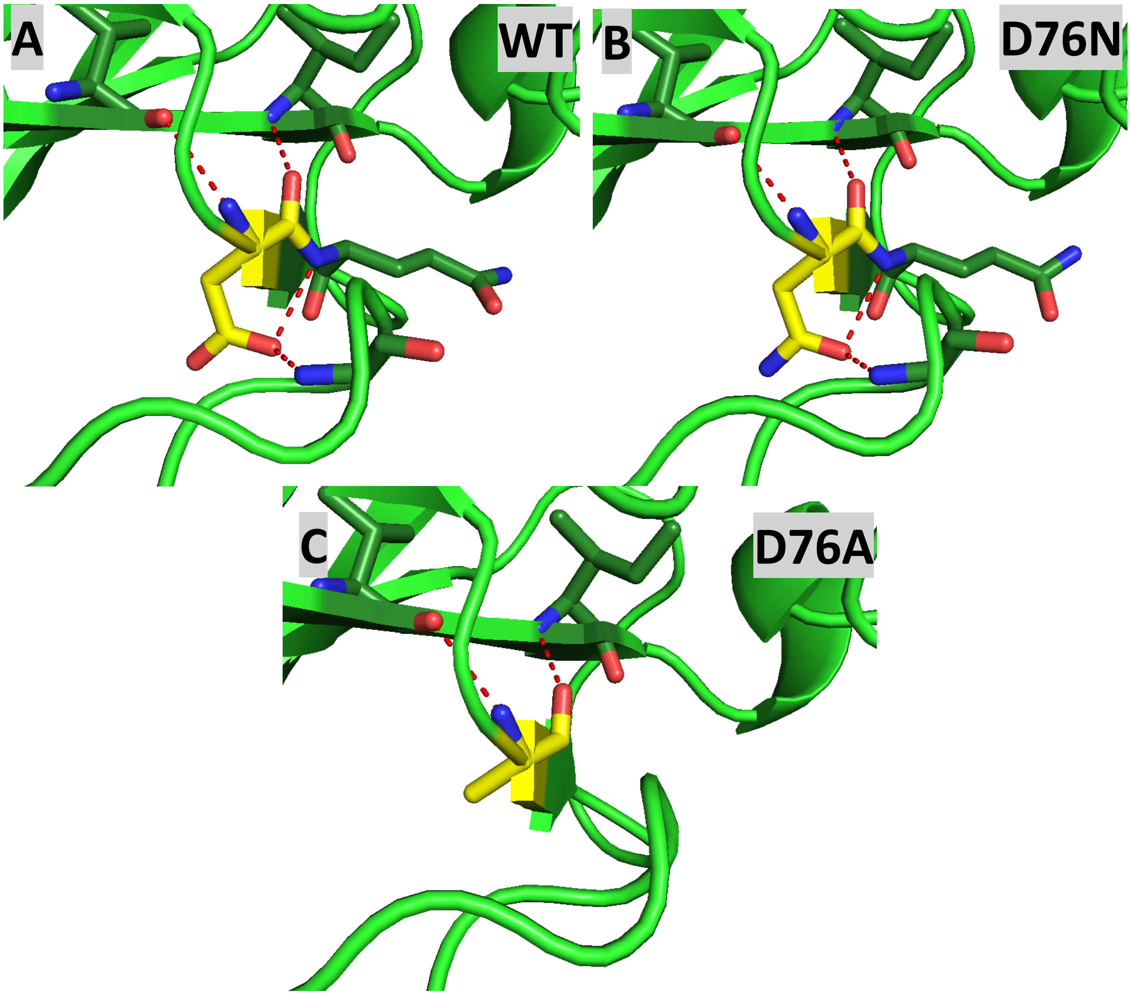


**Figure S18. Predicted hydrogen bonds involving residue 76 in wild-type *Gt*EXPN_133317 (A; Asp76) and its D76N mutant (B) and D76A mutant (C).** The structures were predicted using AlphaFold3 and hydrogen bonds (red dashed lines) were assigned using PyMOL. Carbons in residue 76 are colored yellow. The figure shows that Asp76 in the wild-type makes four hydrogen bonds, two through its main chain and two through its side chain **(A)**. All four bonds involve main chain atoms in the rest of the protein. These bonds are still possible in the D76N mutant, whereas the two side chain-mediated hydrogen bonds, to the main chain amides of Ala30 and Gln77, are no longer possible in the D76A mutant.

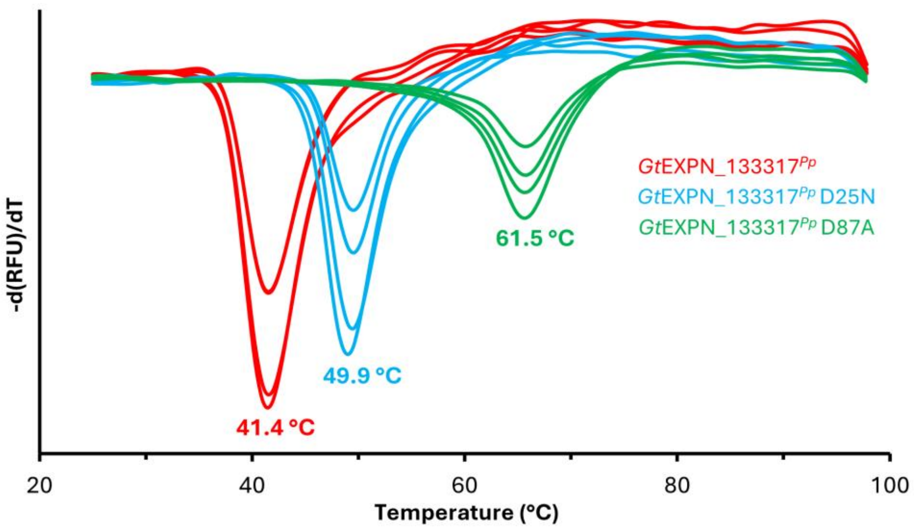


**Figure S19. Thermal stability of *Gt*EXPN_133317*$^{Pp}$* and its D25N and D87A mutants.** The figure shows the melting curves and apparent melting temperatures ($T_{\mathrm{m\ (app)}}$) of *Gt*EXPN_133317$^{Pp}$ wild-type (red lines), *Gt*EXPN_133317$^{Pp}$ D25N (blue lines) and *Gt*EXPN_133317$^{Pp}$ D87A (green lines). The derivative of the fluorescence signal ((-d(RFU)/dT), RFU= relative fluorescence units) is plotted against temperature. 30 µM of protein supplemented with SYPRO orange (fluorescent dye) was heated from 24 °C to 98 °C at 1.42 °C/min. Four replicates were performed. The average $T_{\mathrm{m\ (app)}}$ is indicated for each protein. The standard deviation in the determined $T_m$ values was below ±0.7 °C. In a set of completely independent experiments, and using another protein preparation, the $T_m$ of the wild-type enzyme was determined to be 42.3 °C (**Figure S10**), which is close to the value of 41.4 °C shown here.

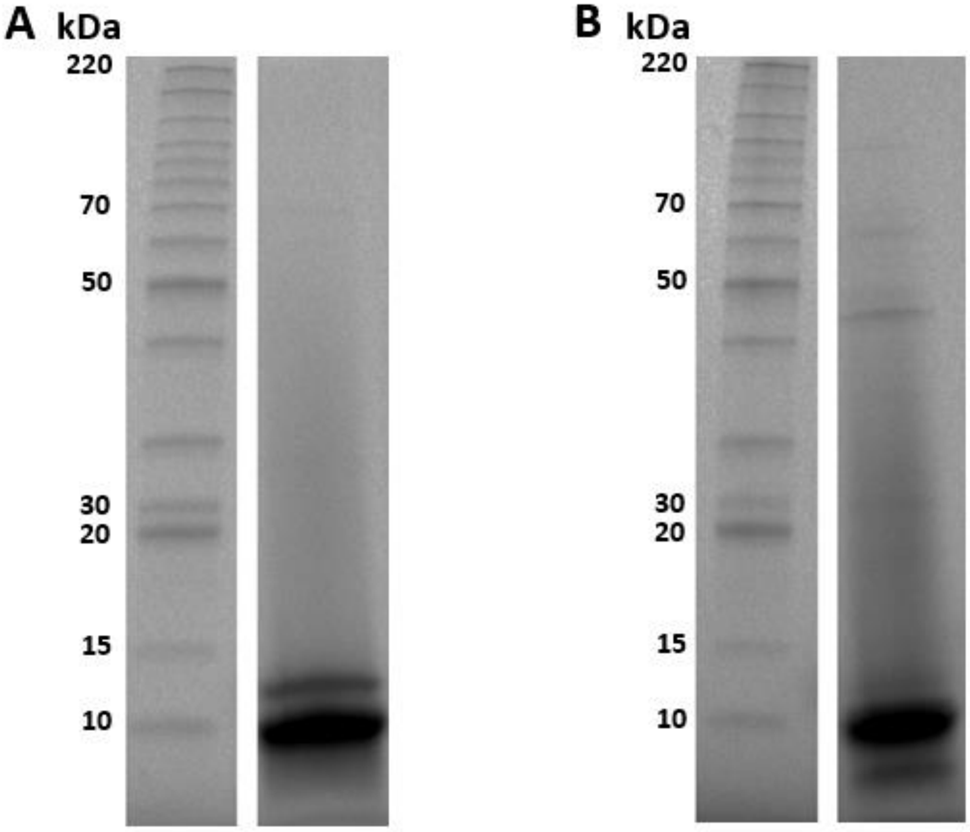


**Figure S20. SDS-PAGE of *Gt*EXPN_133317$^{Pp}$ mutants D87A (A) and D25N (B).** Each gel shows a lane with BenchMark™ Protein Ladder (left) and a sample of purified *Gt*EXPN_133317$^{Pp}$ mutant D87A (50 µg, panel **A**) or D25N (45 µg, panel **B**). The theoretical molecular masses of the protein variants are for (A) *Gt*EXPN_133317$^{Pp}$ D87A, 12417.70 Da and for (B) *Gt*EXPN_133317$^{Pp}$ D25N, 12460.72 Da, as calculated using Expasy's ProtParam tool (https://web.expasy.org/protparam/) and taking into account the formation of two disulfide bridges in the mature proteins. Note that the mutants have the same apparent molecular mass as the *P. pastoris*-produced wild-type protein (**Figure S7A**). The masses of marker proteins are indicated on the left side in kilodaltons (kDa). The gels were cropped to only show relevant gel lanes.

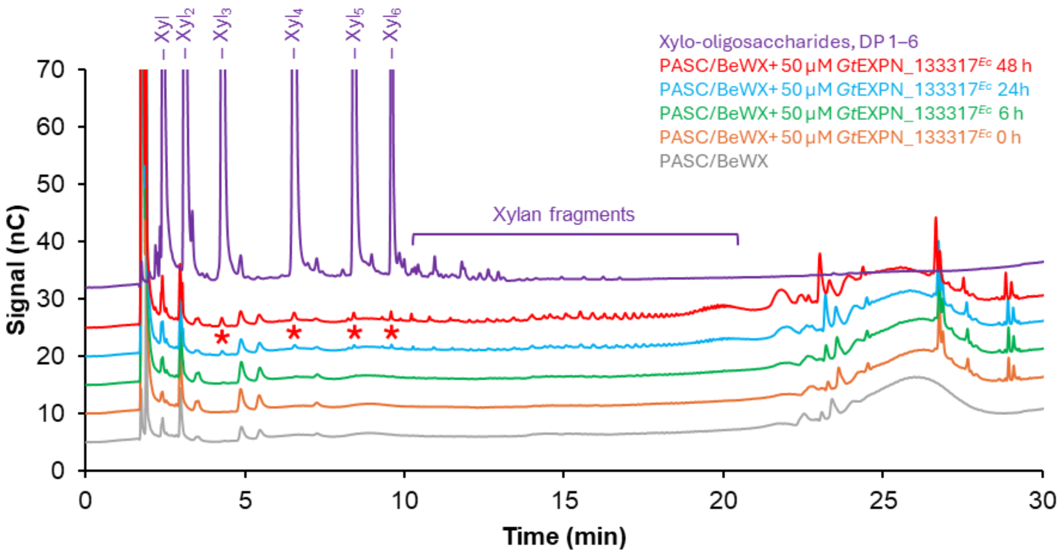


**Figure S21. HPAEC-PAD analysis of products generated over time in a reaction of *Gt*EXPN_133317$^{Ec}$ with a 1:1 (w/w) mixture of PASC and beechwood xylan.** In the reactions, 50 µM *Gt*EXPN_133317$^{Ec}$ was incubated with 0.1% (*w*/*v*) beechwood xylan + 0.1% (*w*/*v*) PASC at 30°C and pH 5.0, for 0 (orange line), 6 (green line), 24 (blue line) and 48 (red line) hours with horizontal shaking at 200 rpm. A 72 h reaction, which is not included in the Figure, showed product levels that were very similar to those observed for the 48 h reaction. The identified linear xylo-oligosaccharides (with DP 2–6) are marked with red asterisks. The chromatogram for a control reaction without enzyme is shown with a grey line. All reactions were performed in triplicates, with similar results; a typical chromatogram is shown. Xylo-oligosaccharides with DP 1–6 (purple), each at 0.025 mg/mL concentration, were used as standard. Note that when zooming in on the y-axis (compared to **Figures 4, 6 and S12**), minor impurities (up to 5%; presumably xylan-derived oligosaccharides) become more visible in the xylo-oligosaccharide standards.

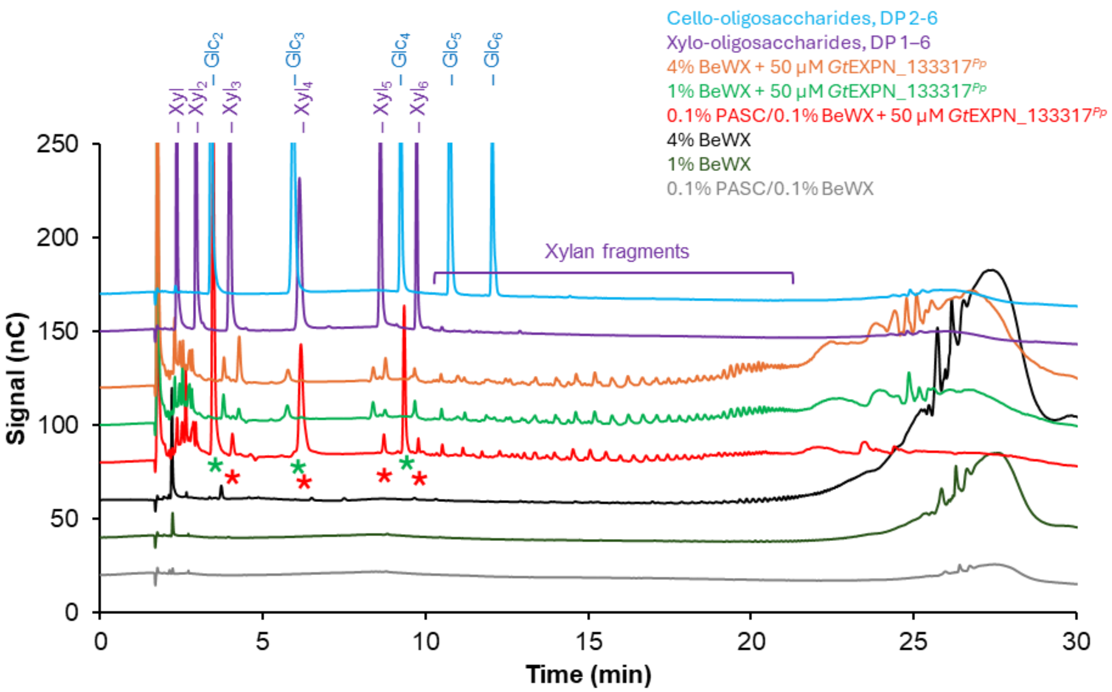


**Figure S22. HPAEC-PAD analysis of products generated over time in a reaction of *Gt*EXPN_133317$^{Pp}$ with a 1:1 (w/w) mixture of 0.1% PASC and 0.1% beechwood xylan and 1% and 4% beechwood xylan.** In the reactions, 50 µM *Gt*EXPN_133317$^{Pp}$ was incubated with 0.1% (*w*/*v*) beechwood xylan and 0.1% (*w*/*v*) PASC (red line), 1% (*w*/*v*) beechwood xylan (green line), or 4% (*w*/*v*) beechwood xylan (orange line) at 37°C and pH 5.0, for 48 h with horizontal shaking at 200 rpm. The identified linear xylo-oligosaccharides [with a degree of polymerization (DP) 3–6] are marked with red asterisks, and the identified cello-oligosaccharides (with DP 2–4) are marked with green asterisks. Control reactions without enzyme are shown with grey [0.1% (*w*/*v*) beechwood xylan and 0.1% (*w*/*v*) PASC], dark green [1% (*w*/*v*) beechwood xylan], and black [4% (*w*/*v*) beechwood xylan] lines. All reactions were performed in triplicates, with similar results; a typical chromatogram is shown. Xylo-oligosaccharides with DP 1–6 (purple line) and cello-oligosaccharides with DP 2–6 (blue line), each at 0.025 mg/mL concentration, were used as standard.

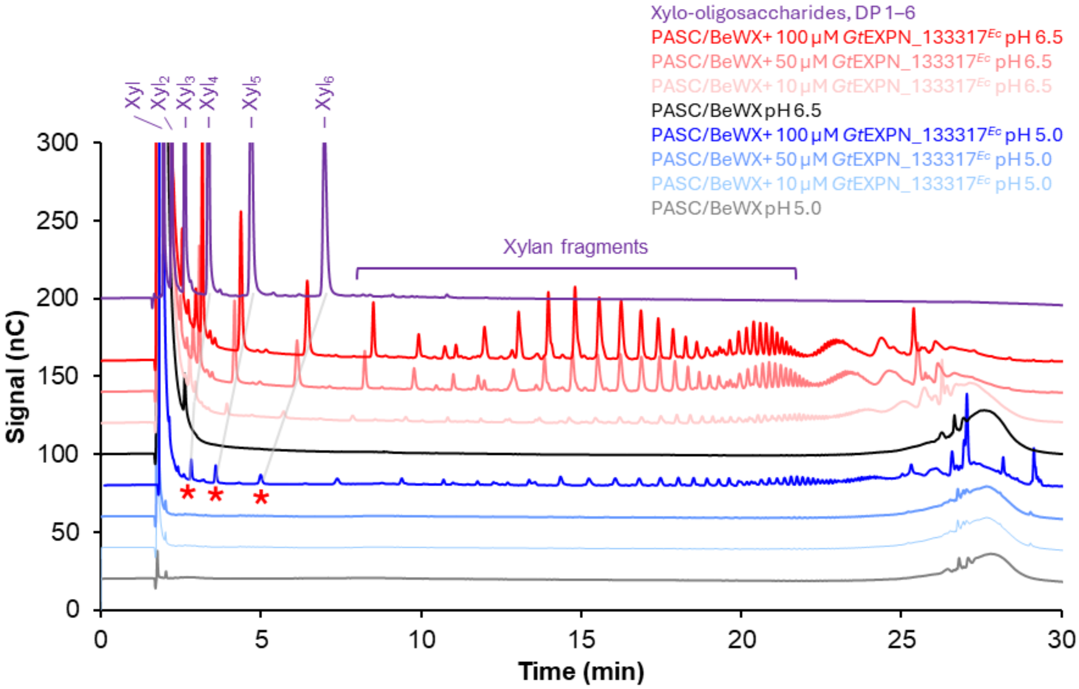


**Figure S23. HPAEC-PAD analysis of products generated over time in a reaction of *Gt*EXPN_133317*^Ec^* with a 1:1 (w/w) mixture of PASC and beechwood xylan at pH 5.0 and pH 6.5.** In the reactions, 10 µM, 50 µM, and 100 µM *Gt*EXPN_133317$^{Ec}$ was incubated with 0.1% (*w*/*v*) beechwood xylan + 0.1% (*w*/*v*) PASC at 30°C for 48 hours with horizontal shaking at 300 rpm. The reactions with 10 µM, 50 µM, and 100 µM of *Gt*EXPN_133317 are shown in light, middle, and dark colors, respectively, and were performed at pH 5.0 (blue lines) or pH 6.5 (red lines). Control reaction without enzyme is shown with dark grey line. All reactions were performed in triplicates, with similar results; a typical chromatogram is shown. Xylo-oligosaccharides with DP 1–6 (purple line), each at 0.025 mg/mL concentration, were used as standard. Despite a shift in the retention time between the chromatograms, the peaks in the chromatograms clearly follow the same pattern. The identified linear xylo-oligosaccharides (with DP 4–6) are marked with red asterisk below the dark blue line, and the corresponding peaks are connected with a light grey guideline.

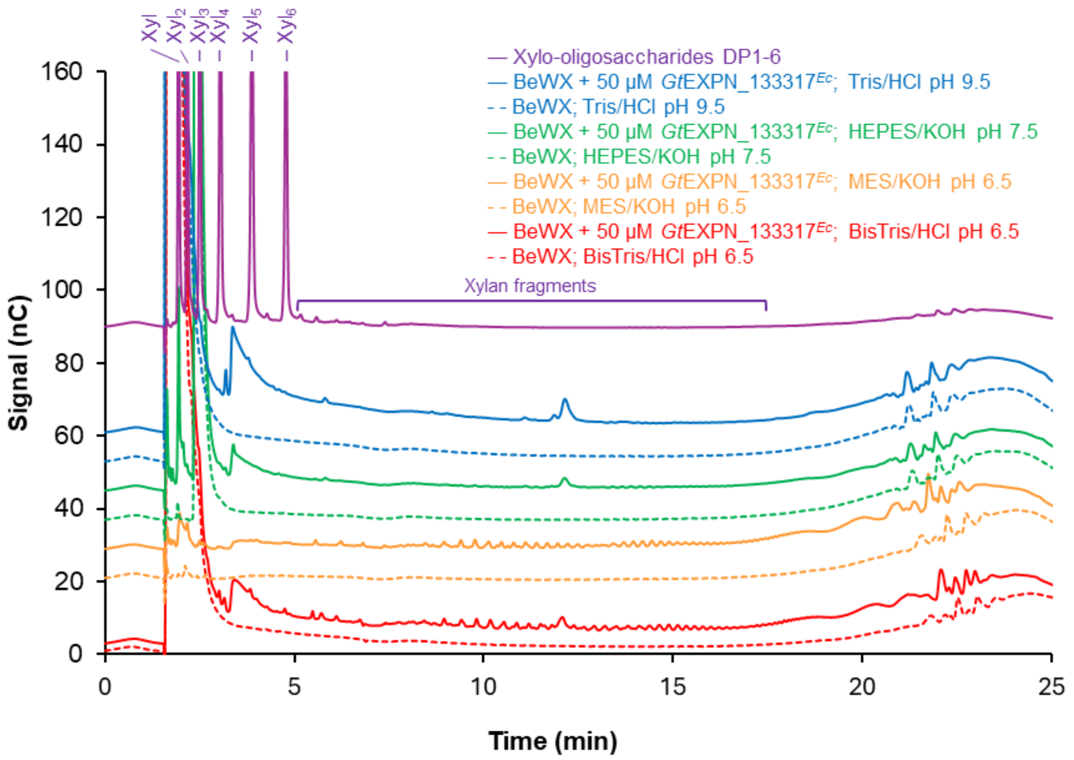


**Figure S24. HPAEC-PAD analysis of products generated in reactions of *Gt*EXPN_133317$^{Ec}$ with 0.2% (*w*/*v*) beechwood xylan at various pH.** In the reactions, 50 µM *Gt*EXPN_133317$^{Ec}$ was incubated with 0.2% (*w*/*v*) beechwood xylan at 30°C for 48 h with horizontal shaking at 160 rpm. The reactions were performed in the following buffers: 50 mM BisTris/HCl pH 6.5 (red line), 25 mM MES/KOH pH 6.5 (orange line), 25 mM HEPES/KOH pH 7.5 (green line) and 25 mM Tris/HCl pH 9.5 (blue line). Corresponding control reactions without enzyme are shown with dashed lines. All reactions were performed in triplicates, with similar results; a typical chromatogram is shown. Xylo-oligosaccharides with DP 1–6 (purple line), each at 0.025 mg/mL concentration, were used as standard.

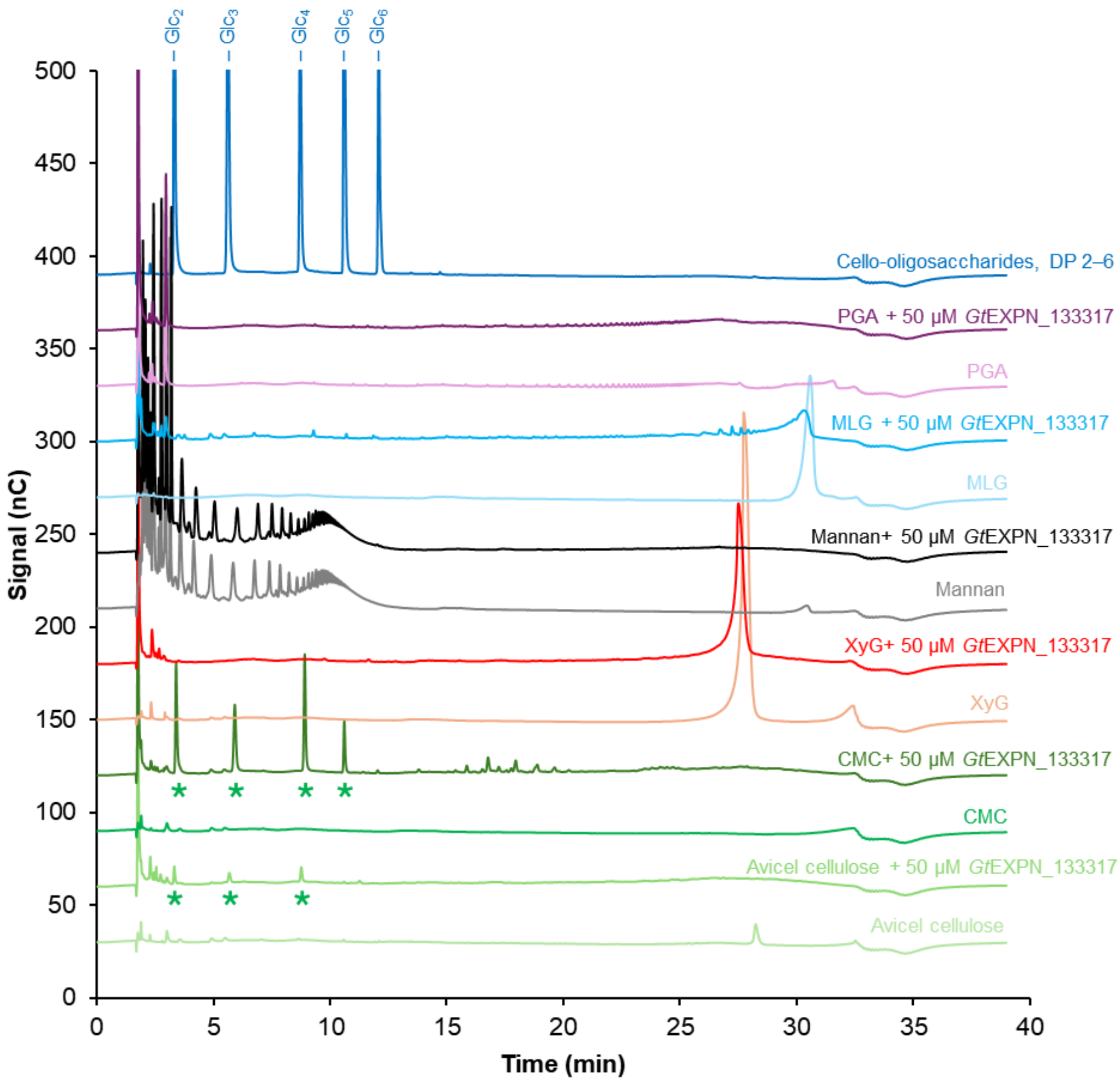


**Figure S25. HPAEC-PAD analysis of products generated over time in reactions with *Gt*EXPN_133317*Pp* and several PCW-derived substrates.** In the reactions, 50 µM *Gt*EXPN_133317*Pp* was incubated with Avicel microcrystalline cellulose (light green line, second from the bottom of the graph), carboxymethylcellulose (CMC; dark green line, fourth from the bottom of the graph), xyloglucan from tamarind seed (XyG; red line), mannan from ivory nut (black line), mixed-linkage β-glucan from barley [β-(1→3),(1→4)-glucan; MLG; blue line], and polygalacturonic acid (PGA; purple line) at 37°C for 48 h with horizontal shaking at 200 rpm. The identified cello-oligosaccharides [with a degree of polymerization (DP) 2–5] are marked with green asterisks. Control reactions without the enzyme are shown with lines below each reaction with enzyme. All reactions were performed in duplicates, with similar results; a typical chromatogram is shown. Cello-oligosaccharides with DP 2–6 (dark blue line) at 0.025 mg/mL concentration were used as standard.

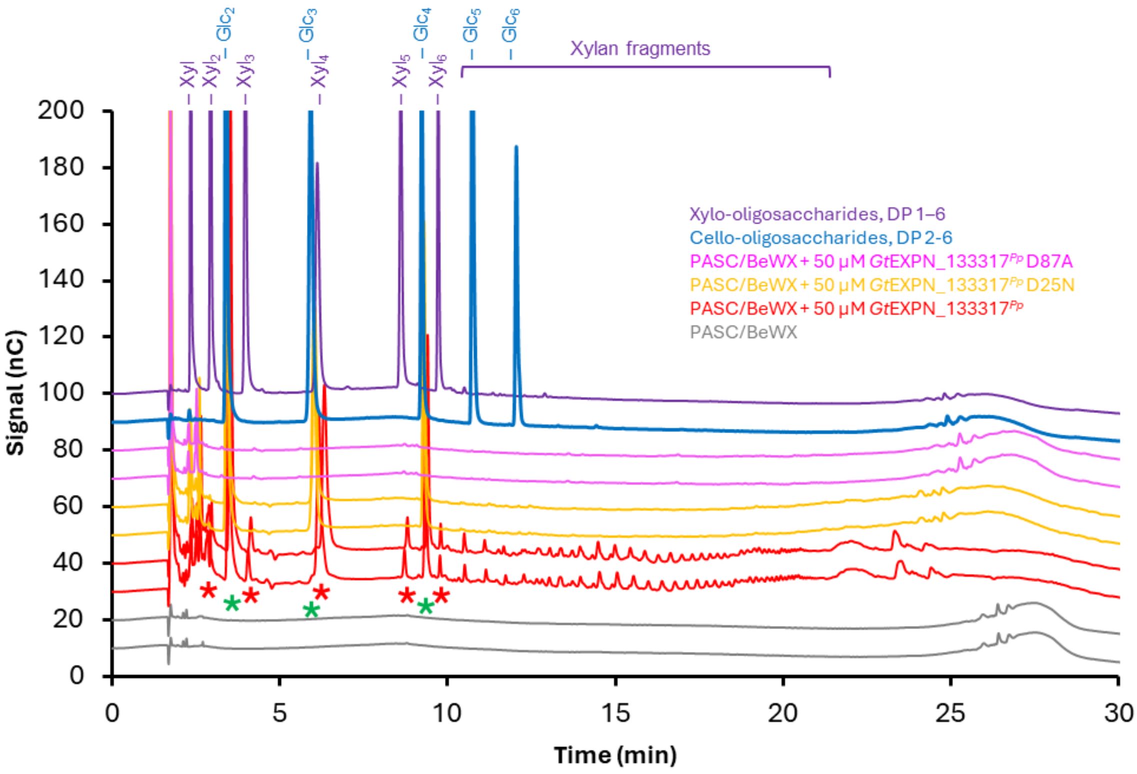


**Figure S26. HPAEC-PAD analysis of products generated in a reaction with *Gt*EXPN_133317$^{Pp}$ or its mutants and a 1:1 (w/w) mixture of PASC and beechwood xylan.** In the reactions, 50 µM *Gt*EXPN_133317$^{Pp}$ (red lines), *Gt*EXPN_133317$^{Pp}$ D25N (orange lines) or *Gt*EXPN_133317$^{Pp}$ D87A (magenta lines) was incubated with 0.1% (*w*/*v*) beechwood xylan + 0.1% (*w*/*v*) PASC at 37°C, pH 5.0, for 48 h with horizontal shaking at 200 rpm. The identified cello-oligosaccharides (with degree of polymerization, DP, 2–4) are marked with green asterisks; the identified linear xylo-oligosaccharides (with DP 2–6) are marked with red asterisks. The result for a control reaction without enzyme is shown with grey lines. All reactions were performed in duplicates, with similar results; the figure shows chromatograms for both reactions. Cello-oligosaccharides with a DP 2–6 (blue line) and xylo-oligosaccharides with DP 1–6 (purple line), each at 0.025 mg/mL concentration, were used as standard.

**Table S1. Proteins with EXPN annotation in the genome of *Gloeophyllum trabeum*.** The annotated genome (13) was accessed at the JGI MycoCosm website (https://mycocosm.jgi.doe.gov/mycocosm/home). Signal peptides (predicted with the SignalP-5.0 server; https://services.healthtech.dtu.dk/services/SignalP-5.0/) are highlighted in grey.

| UniProt ID | FASTA sequence |
|---|---|
| S7QIX4* | >jgi\|Glotr1_1\|34542\|e_gw1.00002.2154.1<br>MFKRSTFTTLVLATSWLPALLTSASVIDPLGSRSAAYARYTTAHSLGERYAFDVRDGWQSINVTDLQYKY<br>TNSGNNLATPKGNILAGASVSHALDSVWNSLKGLGKAEDVTITWYTGHDLENPSCWANSGWAPTDASFAC<br>ALTLEGWTTRPKCFKFLELCNGPKKCTFVRVVDTCAGCAPGSKHVDLTKAAFSQLADVDQGLLTVQMRLA<br>TEPSTW |
| S7Q1Z7 | >jgi\|Glotr1_1\|44870\|e_gw1.00010.546.1<br>MYFSTLTIVSFLSLLVGFTQAAPAAFSLEARNATELAARDSNVRLTWYDVGLGACGKTNKASDYVSPYMS<br>SAEFGSGYPGPHCFKQIKITANGKSATATIVDKCPGCSKGGLDLSKGLFSHFANPDVGVLTGSWNYV |
| S7Q207 | >jgi\|Glotr1_1\|45373\|e_gw1.00010.858.1<br>MYFSILAFVSFLSLIASFAQGAPTEFTLARRNETELARRVSNARLTWYQVGLGACGIANQPSDFVVALDA<br>PDFGSGYPGPHCFQHVSITANGKTATAEIVDKCPGCPEGGLDLSEGLFSFFADPGVGVLTGSWNYV |
| S7RLK0 | >jgi\|Glotr1_1\|130888\|gm1.7195_g<br>MYFSTLAILSFFSFLAGFALAAPTEFALEARNATGLSARDSNARLTWYDVGLGACGKTNKASDYVVALNA<br>AEFGSGYPGPHCFKQIKITANGKSATATIVDQCPGCPKGGLDLSKGLFSHFANPDVGVLTGSWSYL |
| S7Q1Y4 | >jgi\|Glotr1_1\|130892\|gm1.7199_g<br>MYSPSLVLFSFLATLLCLAQAASIHWNSTDVVVPGEKSLAKRVDNAKLTWFSVGLGACGHNNVDSDLVLA<br>LSVADFGAGYPGPNCGRQVRITANGKTATGTVVDKCPGCPAGALDLSLGLFKIFADPAVEVTYGSWTYV |
| S7Q1Y8 | >jgi\|Glotr1_1\|130898\|gm1.7205_g<br>MFISVVPVVSFLAVMAGLAQGSPTEFTAVRRNGTDLDKRVNNARMTYYNVGLGACGITNQPTDFVVALDA<br>TDFGSGYPGPHCFQHISITANGKTATAEIVDKCPGCPVGGLDLSEGLFSFFAPLSTGVIYGSWNYV |
| S7PTB8*** | >jgi\|Glotr1_1\|133317\|gm1.9624_g<br>MVRLALFTTVVASVLAAFASPVPAEDATTLEKRVTHSGRGTWFDVGLGACGEYNVNSDHIVAISAARWGT<br>GANCGQWIHITNTANGKSAYGLTRDQCPGCGVDDLDLSPSLFQELGTLDQGVLSISWHFENKAFSP |
| S7Q2S6** | >jgi\|Glotr1_1\|45028\|e_gw1.00010.655.1<br>NSRSSTATWFTPGLGACGANSLSTDFVVALSPSDYAAGAHCFTHITVNFNGAATDATVFDLCPGCAPGSI<br>DLSPAAFSALASLGAGRVKVDWAFA<br>>Glotr1_1\|45028_corr<br>MFSAVKSLSLALLASLLSAGAVSAASGDATWFTPGLGACGANSLSTDFVVALSPSDYAAGAHCF<br>THITVNFNGAATDATVFDLCPGCAPGSIDLSPAAFSALASLGAGRVKVDWAFA |
| S7Q6B7 | >jgi\|Glotr1_1\|60937\|estExt_Genewise1.C_00007_t10168<br>MMARFSSYFAAALLAFASLLSMVAAVPVEAEHLEARSKTYQGRGTWFNVGLGNCGKENVDSDLIVALDTA<br>TYASGKHCDKYITIVDTKNGKKAKAMVRDSCPSCGVGSIDMSPALFKHFDSLDTGVISVKWWFN |

* Compared to other EXPNs, *Gt*EXPN_34542 has an N-terminal extension of 65 amino acids, for which AlphaFold2 could not predict a structure; these residues are shown in orange. Similar sequences appear in proteins annotated as EXPN in other organisms.

** The N-terminal sequence of *Gt*EXPN_45028, including the signal peptide and the N terminus of the mature protein, is missing from the annotated protein (fragment). Note that a potential start codon can be found 134 bases upstream (towards the 5' end) from the start of the annotated fragment. The 5' sequence of this extended precursor mRNA translates to an N-terminal amino acid sequence that is recognized as a signal peptide and thus may indicate the correct start site for the protein. To ensure that the translated sequence remains "in frame," the presence of a splicing site at the extended 5' region of the precursor mRNA is necessary. This extended 5' region contains a canonical splice site, resulting in a 5′GU…AG3′ intron (13) comprising the nucleotides 86–153 of the extended precursor mRNA. In this table, the corrected N-terminal amino acid sequence is shown in blue; and the correct EXPN sequence is indicated below the currently annotated sequence. Alignment of the corrected protein sequence to the proteins with the highest similarity in the UniProt database indicates that the N terminus with a highly conserved G(D/Q)(A/G)T(W/Y/F)(Y/F) motif is restored after sequence correction.

*** The protein that was studied in detail in this study.

**Table S2A. Transcript levels of *G. trabeum* EXPNs on aspen wood wafers.** In the study by Zhang et al. (14), only eight of nine possible EXPN transcripts were detected. The table shows gene expression levels expressed as reads per kilobase of transcript per million mapped reads (RPKM), with numbers for *Gt*EXPN_130898 and *Gt*EXPN_133317 appearing on a green and yellow background, respectively. Transcriptome analysis was done using RNA-seq on *G. trabeum* growing along the length of wood wafers, with samples taken at set distances behind the hyphal front (0-5 mm, early decay; 15-20 mm, mid decay; 30-35 mm, late decay). The values have been reported previously in Table S3 by Zhang et al. (14).

| UniProt ID | JGI ID (Glotr1_1) | 0-5 mm (RPKM) | 15-20 mm (RPKM) | 30-35 mm (RPKM) | Maximum (RPKM) |
|---|---|---|---|---|---|
| S7QIX4 | 34542 | 3.9788 | 2.9468 | 3.6295 | 3.9788 |
| S7Q1Z7 | 44870 | 7.7404 | 4.6131 | 3.7343 | 7.7404 |
| S7Q207 | 45373 | 2.1058 | 4.0315 | 3.9670 | 4.0315 |
| S7RLK0 | 130888 | 1.2992 | 0.6148 | 0.7213 | 1.2992 |
| S7Q1Y4 | 130892 | 7.8324 | 10.4712 | 9.1336 | 10.4712 |
| S7Q1Y8 | 130898 | 13.6943 | 15.7055 | 17.4724 | 17.4724 |
| S7PTB8 | 133317 | 749.9677 | 1424.7250 | 697.4629 | 1424.7250 |
| S7Q2S6 | 45028 | 0.0000 | 0.0000 | 0.0000 | 0.0000 |
| S7Q6B7 | 60937 | 9.5256 | 7.8665 | 9.8178 | 9.8178 |

**Table S2B. Transcript levels of *G. trabeum* EXPNs on Japanese cedar, cellulose, and glucose.** In the study by Umezawa et al. (15) all nine possible EXPN transcripts were detected. The table shows gene expression levels expressed as transcripts per million (TPM) mapped reads, with numbers for *Gt*EXPN_130898 and *Gt*EXPN_133317 appearing on a green and yellow background, respectively. Transcriptome analysis was done using RNA-seq for *G. trabeum* growing in liquid media containing glucose, Avicel, or conifer wood from Japanese cedar. The values (averages of three biological replicates) have been reported previously in Table S2 by Umezawa et al. (15).

| Protein ID | JGI ID (Glotr1_1) | Glucose, average (TPM) | Avicel, average (TPM) | Cedar, average (TPM) |
|---|---|---|---|---|
| S7QIX4 | 34542 | 0.8934 | 0.7966 | 1.2139 |
| S7Q1Z7 | 44870 | 16.7435 | 15.8162 | 13.4203 |
| S7Q207 | 45373 | 3.1130 | 10.5177 | 7.4040 |
| S7RLK0 | 130888 | 14.1106 | 5.1557 | 4.8642 |
| S7Q1Y4 | 130892 | 1.1410 | 2.4572 | 2.0275 |
| S7Q1Y8 | 130898 | 15.1556 | 49956.2165 | 14113.2034 |
| S7PTB8 | 133317 | 2367.7828 | 743.8119 | 54.4055 |
| S7Q2S6 | 45028 | 689.7807 | 77.1992 | 41.3293 |
| S7Q6B7 | 60937 | 69.0475 | 5.1749 | 9.4010 |

**Table S3A. Secretome levels of *G. trabeum* EXPNs during growth on spruce wood wafers.** In the study by Presley et al. (16), from which the data shown is taken, peptides of only two of the nine EXPN proteins were detected. Secretome analysis was done by tryptic digestion of protein extracts from spruce wafers degraded by *G. trabeum*, dividing the wood wafers along the length at set distances behind the hyphal front (0-5 mm, early decay; 10-15 mm, mid decay; 20-25 mm, late decay), with subsequent LC-$MS^2$ analysis. The table shows the number of $MS^2$ spectra assigned to each protein, with numbers for *Gt*EXPN_44870 and *Gt*EXPN_133317 appearing on orange and yellow backgrounds, respectively. These values have been reported previously in Dataset S1 by Presley et al. (16). Abbreviations: b.i.l., below identification limit (two observed peptides/protein in at least one of the samples; proteins that were never detected get "b.i.l.", whereas proteins that were detected in some samples get 0 for samples in which they were not detected).

| UniProt ID | JGI ID (Glotr1_1) | 0-5 mm | 10-15 mm | 20-25 mm |
|---|---|---|---|---|
| S7QIX4 | 34542 | b.i.l. | b.i.l. | b.i.l. |
| S7Q1Z7 | 44870 | 2 | 0 | 0 |
| S7Q207 | 45373 | b.i.l. | b.i.l. | b.i.l. |
| S7RLK0 | 130888 | b.i.l. | b.i.l. | b.i.l. |
| S7Q1Y4 | 130892 | b.i.l. | b.i.l. | b.i.l. |
| S7Q1Y8 | 130898 | b.i.l. | b.i.l. | b.i.l. |
| S7PTB8 | 133317 | 2 | 3 | 0 |
| S7Q2S6 | 45028 | b.i.l. | b.i.l. | b.i.l. |
| S7Q6B7 | 60937 | b.i.l. | b.i.l. | b.i.l. |

**Table S3B. Secretome levels of *G. trabeum* EXPNs during growth on aspen wood wafers.** In a study by Presley et al. (17), from which the data shown is taken, peptides of three of the nine possible EXPN proteins were detected. Secretome analysis was done by tryptic digestion of protein extracts from spruce wafers degraded by *G. trabeum*, dividing the wood wafers along the length at set distances behind the hyphal front (0-5 mm, early decay; 15-20 mm, mid decay; 30-35 mm, late decay), with subsequent LC-$MS^2$ analysis. The table shows the number of $MS^2$ spectra assigned to each protein, with numbers for *Gt*EXPN_44870 and *Gt*EXPN_133317 appearing on orange and yellow backgrounds, respectively. The values have been reported previously in Dataset S1 by Presley et al. (17). Abbreviation: b.i.l., below identification limit (two observed peptides/protein in at least one of the samples; proteins that were never detected get "b.i.l.", whereas proteins that were detected in some samples get 0 for samples in which they were not detected).

| UniProt ID | JGI ID (Glotr1_1) | 0-5 mm (2 replicates) | | 15-20 mm (3 replicates) | | | 30-35 mm (3 replicates) | | |
|---|---|---|---|---|---|---|---|---|---|
| S7QIX4 | 34542 | b.i.l. | | b.i.l. | | | b.i.l. | | |
| S7Q1Z7 | 44870 | 4 | 12 | 12 | 12 | 0 | 0 | 0 | 0 |
| S7Q207 | 45373 | b.i.l. | | b.i.l. | | | b.i.l. | | |
| S7RLK0 | 130888 | 0 | 0 | 0 | 12 | 0 | 0 | 0 | 0 |
| S7Q1Y4 | 130892 | b.i.l. | | b.i.l. | | | b.i.l. | | |
| S7Q1Y8 | 130898 | b.i.l. | | b.i.l. | | | b.i.l. | | |
| S7PTB8 | 133317 | 3 | 24 | 0 | 0 | 0 | 0 | 0 | 0 |
| S7Q2S6 | 45028 | b.i.l. | | b.i.l. | | | b.i.l. | | |
| S7Q6B7 | 60937 | b.i.l. | | b.i.l. | | | b.i.l. | | |

**Table S4. Protein variants identified in the purified *Gt*EXPN_133317$^{Pp}$ preparation.** The theoretical molecular masses were calculated using Expasy's ProtParam tool (https://web.expasy.org/protparam/), taking into account the formation of two disulfide bridges in the mature protein; the *m/z* values were obtained using MALDI-ToF MS (see **Figure S8B**). There is a consistent 10-11 Da difference ('Difference' column) between the theoretical molecular mass and the observed protein mass (with a charge of *z*=1). The sequence of the full-length protein is shown for reference. Despite the unknown deviation of 10 Da, this MALDI-TOF MS analysis of whole proteins clearly shows that the proteins are not glycosylated. Tryptic digestion and LC-MS analysis of *Gt*EXPN_133317$^{Pp}$ (**Dataset S1**) did not reveal clear signs of post-translational modifications either and confirmed the identity of the protein.

| Protein sequence | Theoretical molecular mass (Da) | *m/z* | Difference |
|---|---|---|---|
| **S1–P117 (Full-length):**<br>SPVPAEDATT LEKRVTHSGR GTWFDVGLGA<br>CGEYNVNSDH IVAISAARWG TGANCGQWIH<br>ITNTANGKSA YGLTRDQCPG CGVDDLDLSP<br>SLFQELGTLD QGVLSISWHF ENKAFSP | 12461.71 | Not observed | - |
| **E12–P117:**<br>EKRVTHSGR GTWFDVGLGA<br>CGEYNVNSDH IVAISAARWG TGANCGQWIH<br>ITNTANGKSA YGLTRDQCPG CGVDDLDLSP<br>SLFQELGTLD QGVLSISWHF ENKAFSP | 11379.53 | 11390.10 | +10.6 |
| **K13–P117:**<br>KRVTHSGR GTWFDVGLGA<br>CGEYNVNSDH IVAISAARWG TGANCGQWIH<br>ITNTANGKSA YGLTRDQCPG CGVDDLDLSP<br>SLFQELGTLD QGVLSISWHF ENKAFSP | 11250.42 | 11261.03 | +10.6 |
| **V15–P117:**<br>VTHSGR GTWFDVGLGA<br>CGEYNVNSDH IVAISAARWG TGANCGQWIH<br>ITNTANGKSA YGLTRDQCPG CGVDDLDLSP<br>SLFQELGTLD QGVLSISWHF ENKAFSP | 10966.05 | 10976.79 | +10.7 |
| **V15–A114:**<br>VTHSGR GTWFDVGLGA<br>CGEYNVNSDH IVAISAARWG TGANCGQWIH<br>ITNTANGKSA YGLTRDQCPG CGVDDLDLSP<br>SLFQELGTLD QGVLSISWHF ENKA | 10634.67 | 10644.63 | +10.0 |

**Table S5. Theoretical *m/z* values for singly charged glucuronoxylan-derived compounds.** The listed *m/z* values correspond to the $Na^+$-adducts of non-glucuronylated ($Xyl_n$), singly 4-*O*-methylglucuronylated ($Xyl_n$MeGlcA) and doubly 4-*O*-methylglucuronylated [$Xyl_n$(MeGlcA)$_2$] compounds, where *n* is the number of xylosyl units in the backbone. The *m/z* values for the compounds where the carboxyl of one 4-*O*-methylglucuronyl substitution forms a $Na^+$-salt (MeGlcA–Na) are also shown. These masses correspond to hydrolysis products. Oxidative cleavage would lead to the formation of compounds with Δ(*m/z*) = +16.00 (hydrated form) or Δ(*m/z*) = −2.02 (anhydro form) compared to the native, i.e., non-oxidized, compounds. Only compounds corresponding to hydrolysis products were detected among the reaction products generated by *Gt*EXPN_133317 acting on glucuronoxylan.

| ***n*** | ***m/z*** **$Xyl_n/Na^+$** | ***m/z*** **$Xyl_n$MeGlcA/$Na^+$** | ***m/z*** **$Xyl_n$(MeGlcA–Na)/$Na^+$** | ***m/z*** **$Xyl_n$(MeGlcA)$_2$/$Na^+$** | ***m/z*** **$Xyl_n$MeGlcA(MeGlcA–Na)/$Na^+$** |
|---|---|---|---|---|---|
| 4 | 569.46 | 759.61 | 781.60 | — | — |
| 5 | 701.58 | 891.73 | 913.71 | — | — |
| 6 | 833.69 | 1023.84 | 1045.83 | — | — |
| 7 | 965.81 | 1155.96 | 1177.94 | 1346.11 | 1368.09 |
| 8 | 1097.92 | 1288.07 | 1310.05 | 1478.22 | 1500.21 |
| 9 | 1230.04 | 1420.19 | 1442.17 | 1610.34 | 1632.32 |
| 10 | 1362.15 | 1552.30 | 1574.28 | 1742.45 | 1764.43 |
| 11 | 1494.27 | 1684.42 | 1706.40 | 1874.57 | 1896.55 |
| 12 | 1626.38 | 1816.53 | 1838.51 | 2006.68 | 2028.66 |
| 13 | 1758.50 | 1948.65 | 1970.63 | 2138.80 | 2160.78 |
| 14 | 1890.61 | 2080.76 | 2102.74 | 2270.91 | 2292.89 |
| 15 | 2022.72 | 2212.88 | 2234.86 | 2403.03 | 2425.01 |
| 16 | 2154.84 | 2344.99 | 2366.97 | 2535.14 | 2557.12 |
| 17 | 2286.95 | 2477.10 | 2499.09 | 2667.25 | 2689.24 |

**Table S6. Yields of purified *Gt*EXPN_133317 variants**. Abbreviations: *Ec*, produced in *E. coli*; *Pp*, produced in *P. pastoris*; WT, wild-type.

| Protein | Yield (mg protein/L culture) |
|---|---|
| *Gt*EXPN_133317$^{Pp}$ WT | 30 |
| *Gt*EXPN_133317$^{Pp}$ D87A | 30 |
| *Gt*EXPN_133317$^{Pp}$ D25N | 27 |
| *Gt*EXPN_133317$^{Ec}$ WT | 5.5 |

**SI References**


1. F. Armougom *et al.*, Expresso: automatic incorporation of structural information in multiple sequence alignments using 3D-Coffee. *Nucleic Acids Res.* **34**, W604-W608 (2006).
2. G. Courtade, S. B. Le, G. I. Sætrom, T. Brautaset, F. L. Aachmann, A novel expression system for lytic polysaccharide monooxygenases. *Carbohydr. Res.* **448**, 212-219 (2017).
3. C. Manoil, J. Beckwith, A genetic approach to analyzing membrane protein topology. *Science* **233**, 1403-1408 (1986).
4. M. M. Bradford, A rapid and sensitive method for the quantitation of microgram quantities of protein utilizing the principle of protein-dye binding. *Anal. Biochem.* **72**, 248-254 (1976).
5. T. M. Wood, "Preparation of crystalline, amorphous, and dyed cellulase substrates" in Methods in Enzymology. (Academic Press, 1988), vol. 160, pp. 19-25.
6. M. Jensen *et al.*, Engineering chitinolytic activity into a cellulose-active lytic polysaccharide monooxygenase provides insights into substrate specificity. *J. Biol. Chem.* **294**, jbc.RA119.010056 (2019).
7. G. L. Miller, Use of dinitrosalicylic acid reagent for determination of reducing sugar. *Anal. Chem.* **31**, 426-428 (1959).
8. B. Westereng, M. Arntzen, J. W. Agger, G. Vaaje-Kolstad, V. G. H. Eijsink, Analyzing activities of lytic polysaccharide monooxygenases by liquid chromatography and mass spectrometry. *Methods in Molecular Biology* **1588**, 71-92 (2017).
9. H. Østby, J.-K. Jameson, T. Costa, V. G. H. Eijsink, M. Ø. Arntzen, Chromatographic analysis of oxidized cello-oligomers generated by lytic polysaccharide monooxygenases using dual electrolytic eluent generation. *Journal of Chromatography A* **1662**, 462691 (2022).
10. O. A. Hegnar *et al.*, Quantifying oxidation of cellulose-associated glucuronoxylan by two lytic polysaccharide monooxygenases from *Neurospora crassa*. *Appl. Environ. Microbiol.* **87**, e0165221 (2021).
11. J. W. Agger *et al.*, Discovery of LPMO activity on hemicelluloses shows the importance of oxidative processes in plant cell wall degradation. *Proceedings of the National Academy of Sciences* **111**, 6287-6292 (2014).
12. T. R. Tuveng *et al.*, Proteomic investigation of the secretome of *Cellvibrio japonicus* during growth on chitin. *Proteomics* **16**, 1904-1914 (2016).
13. D. Floudas *et al.*, The paleozoic origin of enzymatic lignin decomposition reconstructed from 31 fungal genomes. *Science* **336**, 1715-1719 (2012).
14. J. Zhang, A. T. Silverstein Kevin, D. Castaño Jesus, M. Figueroa, J. S. Schilling, Gene regulation shifts shed light on fungal adaption in plant biomass decomposers. *mBio* **10**, 10.1128/mbio.02176-02119 (2019).
15. K. Umezawa, M. Niikura, Y. Kojima, B. Goodell, M. Yoshida, Transcriptome analysis of the brown rot fungus *Gloeophyllum trabeum* during lignocellulose degradation. *PLOS ONE* **15**, e0243984 (2020).

16. G. N. Presley, J. S. Schilling, Distinct growth and secretome strategies for two taxonomically divergent brown rot fungi. *Appl. Environ. Microbiol.* **83**, e02987-02916 (2017).
17. G. N. Presley, E. Panisko, S. O. Purvine, J. S. Schilling, Coupling secretomics with enzyme activities to compare the temporal processes of wood metabolism among white and brown rot fungi. *Appl. Environ. Microbiol.* **84**, e00159-00118 (2018).
